# Human cytomegalovirus tropism modulator UL148 interacts with SEL1L, a cellular factor that governs ER-associated degradation of the viral envelope glycoprotein, gO

**DOI:** 10.1101/304394

**Authors:** Christopher C. Nguyen, Mohammed N. Siddiquey, Hongbo Zhang, Jeremy P. Kamil

## Abstract

UL148 is a viral endoplasmic reticulum (ER)-resident glycoprotein that contributes to human cytomegalovirus (HCMV) cell tropism. The influence of UL148 on tropism correlates with its potential to promote the expression of glycoprotein O (gO), a viral envelope glycoprotein that participates in a heterotrimeric complex with glycoproteins H and L that is required for infectivity. In an effort to gain insight into mechanism, we used mass spectrometry to identify proteins that co-immunoprecipitate from infected cells with UL148. This approach led us to identify an interaction between UL148 and SEL1L, a factor that plays key roles in ER-associated degradation (ERAD). In pulse-chase experiments, gO was less stable in cells infected with a *UL148*-null mutant HCMV than during wild-type infection, suggesting a potential functional relevance for the interaction with SEL1L. To investigate whether UL148 regulates gO abundance by influencing ERAD, siRNA silencing of either SEL1L or its partner, Hrd1, was carried out in the context of infection. Knockdown of these ERAD factors strongly enhanced levels of gO, but not other viral glycoproteins, and the effect was amplified in the presence of UL148. Furthermore, pharmacological inhibition of ERAD showed similar results. Silencing of SEL1L during infection also stabilized an interaction of gO with the ER lectin OS-9, which likewise suggests that gO is an ERAD substrate. Taken together, our results identify an intriguing interaction of UL148 with the ERAD machinery, and demonstrate that gO behaves as a constitutive ERAD substrate during infection. These findings have implications for understanding the regulation of HCMV cell tropism.

## IMPORTANCE

Viral glycoproteins in large part determine the cell types that an enveloped virus can infect, and hence play crucial roles in transmission and pathogenesis. The glycoprotein H/L heterodimer (gHgL) is part of the conserved membrane fusion machinery that all herpesviruses use to enter cells. In human cytomegalovirus, gHgL participates in alternative complexes in virions, one of which is a trimer of gHgL with glycoprotein O (gO). Here, we show that gO is constitutively degraded during infection by the endoplasmic reticulum-associated degradation (ERAD) pathway, and that UL148, a viral factor that regulates HCMV cell tropism, interacts with the ERAD machinery and slows gO decay. Since gO is required for cell-free virus to enter new host cells, but dispensable for cell-associated spread that resists antibody neutralization, our findings imply that the post-translational instability of a viral glycoprotein provides a basis for viral mechanisms to modulate tropism and spread.

## INTRODUCTION

Human cytomegalovirus (HCMV) is a β-herpesvirus that establishes life-long infection in the human host and causes significant morbidity and mortality in immunocompromised patients. The virus exhibits a remarkably broad cell tropism, infecting a wide array of cell types that include epithelial cells, endothelial cells, fibroblasts, smooth muscle cells, hepatocytes, leukocytes, and hematopoietic cells [reviewed in (1, 2)]. Primary infection is thought to commence with viral replication in mucosal epithelia. The virus then transfers to circulating leukocytes to establish latent infection in hematopoietic progenitor cells (3–5). For horizontal transmission, HCMV productively infects the epithelium of several secretory tissues, allowing for the release of virus into bodily fluids including saliva, breastmilk, and urine (2, 6).

HCMV expresses two alternative envelope glycoprotein H / glycoprotein L (gHgL) complexes that impact its cell tropism: gHgLgO (“trimer”) and gHgL/UL128-UL130-UL131 (“pentamer”) [reviewed in (7)]. Although the pentamer is dispensable for infection of fibroblasts, it is required to infect epithelial cells, endothelial cells, and leukocytes (8–10). Recently, Murrell *et al*. showed that the pentamer promotes cell-to-cell spread that is resistant to antibody neutralization (11). On the other hand, the trimer appears to be essential for the infectivity of cell-free HCMV virions, since *gO*-null mutant viruses are defective for entry into all cell types (12, 13). While the platelet-derived growth factor receptor alpha has been identified as a cellular receptor for the trimer that is required for infection of fibroblasts (14–16), pentamer receptor(s) remain to be demonstrated. Evidence from the murine cytomegalovirus (MCMV) *in vivo* infection model suggests that the MCMV ortholog of the HCMV trimer (gHgL with m74) promotes initial infection of tissues via a cell-free route but is dispensable for subsequent intratissue spread (17). While these and other studies have significantly advanced the understanding of HCMV envelope glycoprotein complexes involved in cell-type specific entry, the mechanisms driving production of virions with distinct glycoprotein profiles remain unclear, despite previous reports that imply that such mechanisms exist (18, 19).

In 2015, we reported the identification of a viral regulator of HCMV envelope gH/gL complexes encoded by the *UL148* gene (20). Disruption of *UL148* in strain TB40/E led to markedly enhanced tropism for epithelial cells accompanied by reduced levels of trimer on virions, and an overall reduction in virion levels of gHgL. These findings, together with results from two studies by Zhou et al. (21, 22), imply that HCMV cell tropism might be influenced by the ratio of pentamer-to-trimer in the viral envelope, and/or by the overall amount of gH complexes in virions. Although our previous work suggested that UL148 increases the amount of trimer that matures beyond the endoplasmic reticulum (ER) to the post-Golgi compartments where virions acquire their infectious envelopes, the molecular details have remained unknown. UL148 localizes exclusively to the ER, and *UL74* (gO) transcript levels were unaffected by disruption of *UL148* (20). Hence, it seems likely that UL148 acts within the ER to either stabilize gO or promote assembly of the trimer. To gain more detailed information concerning potential mechanisms, we embarked to identify proteins that interact with UL148 during infection. Here, we show that UL148 interacts with SEL1L, a core element of the cellular ER-associated degradation (ERAD) pathway. Furthermore, results of experiments carried out to address the functional relevance of the interaction reveal that gO behaves as a constitutive ERAD substrate during HCMV infection.

## RESULTS

### HCMV tropism factor UL148 physically associates with the ERAD adapter SEL1L during infection

To investigate the mechanism by which UL148 promotes gO expression, we sought to identify its interaction partners. To this end, we infected primary human foreskin fibroblasts (HFF) with an recombinant HCMV strain TB40/E that expresses an HA-epitope tagged UL148 (TB_148^HA^). A parallel infection was conducted using a control virus, TB_16^HA^, which expresses HA-tagged UL16. Like UL148, UL16 is an ER resident glycoprotein with type I transmembrane topology (23). Infected cells were harvested for anti-HA immunoprecipitation (IP) at 72 h post infection (hpi). After resolving eluates by SDS-PAGE, we excised silver-stained bands for protein identification by mass spectrometry (MS) analysis (Fig. 1A). The full list of cellular proteins identified from IPs of TB_148^HA^ infections in two independent experiments, but not from IPs of TB_16^HA^, is shown in Table S1. We were intrigued to observe peptides that mapped to factors in the ERAD pathway, including SEL1L, XTP3-B, and OS-9.

**FIG. 1.**
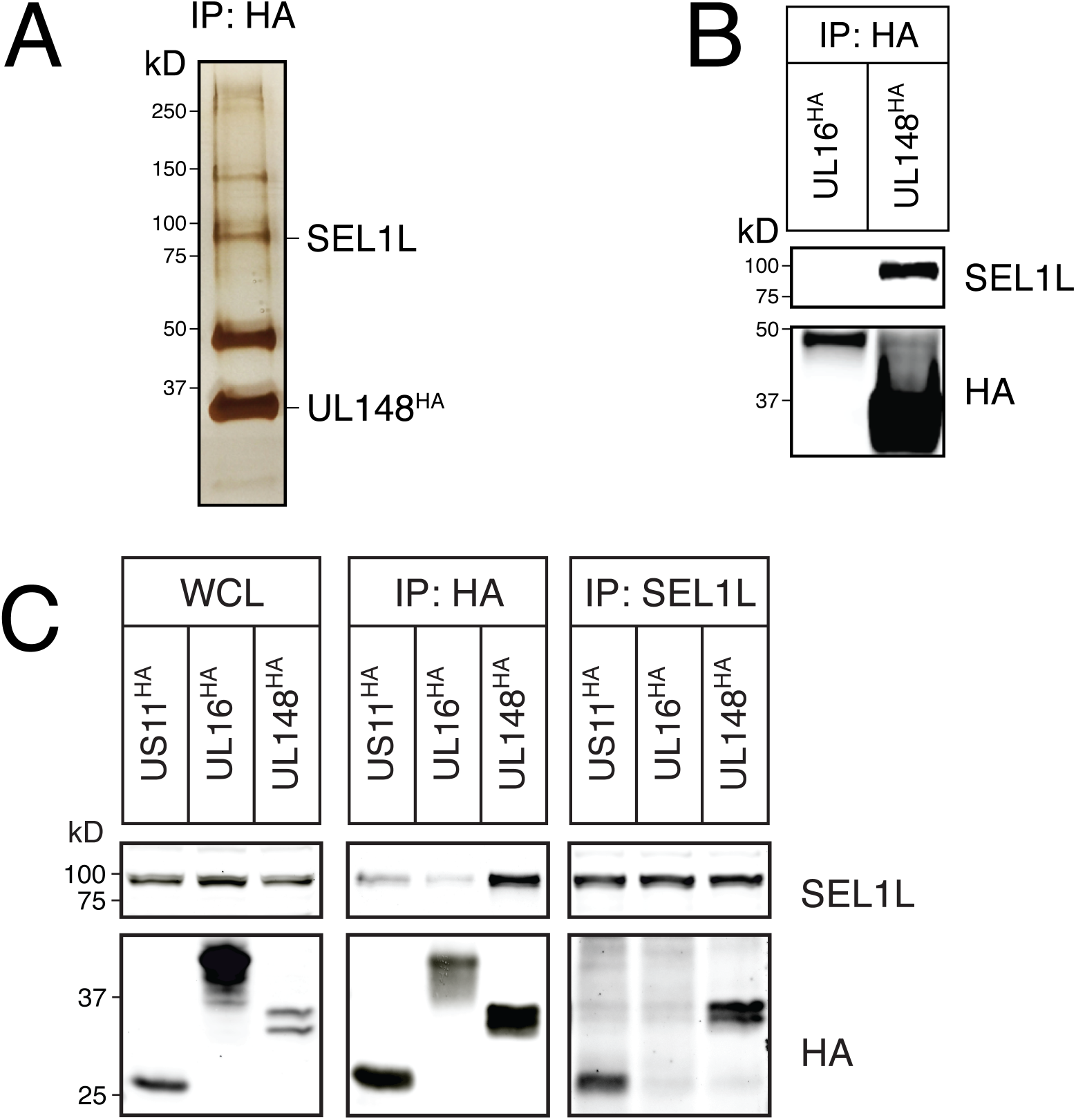
Viral tropism factor UL148 associates with ERAD adapter SEL1L during infection. (A) Silver stain. Fibroblasts (HFF) infected with TB_148HA were lysed at 72 hpi and subjected to HA-IP. Visible silver-stained bands were excised and analyzed by mass spectrometry. The putative association between UL148 and SEL1L was confirmed by HA-IP and western blot of UL148HA complexes from (B) infected cells or (C) 293T cells transfected with plasmids expressing the indicated HCMV ER-resident proteins.

Although co-IP experiments failed to validate several of the putative hits (not shown), we found evidence that suggested a physical association between UL148 and SEL1L (Fig. 1). Results of co-IP experiments from HCMV infected fibroblasts (Fig. 1B), and from plasmid transfected 293T cells (Fig. 1C) supported the possibility that UL148 associates with endogenous SEL1L. HCMV US11, a viral immune-evasin that was included as a positive control known to interact with SEL1L (24), was readily detected in anti-SEL1L IPs, as was UL148. However, detection of SEL1L was less pronounced following IP of HA-tagged US11 than following IP of HA-tagged UL148, which may indicate a more robust interaction with UL148. UL16 and SEL1L were found to co-IP each other poorly or undetectably, consistent with the negative result from infected cells (Fig. 1B-1C). From these results, we concluded that UL148 may physically associate with SEL1L, or with a protein complex that contains it.

### Pulse-chase analysis of gO during WT-and *UL148*-null HCMV infection

SEL1L plays key roles in the degradation of misfolded ER proteins [reviewed in (25, 26)]. In the canonical ERAD pathway, SEL1L acts as an adapter between the ER lectins OS-9 and XTP3-B and the Hrd1 complex (27), of which SEL1L is a stable component (28, 29). The ER lectins are posited to deliver terminally misfolded glycoproteins to the SEL1L-Hrd1 complex, which mediates ubiquitination and translocation of ERAD substrates to the cytosol for proteasomal degradation (27). Given that UL148 is required for high level expression of gO during infection (20) and appears to physically associate with SEL1L (Fig. 1), we sought to determine (i) whether the ERAD pathway targets gO for degradation and (ii) whether UL148 limits degradation of gO.

We therefore carried out pulse-chase experiments to compare gO stability between WT and *UL148*-null HCMV infected cells, using recombinant viruses that express an S-peptide tag at the C-terminus of gO in a manner that avoids disruption of the overlapping *UL73* gene encoding gN (Fig. S1A). Importantly, the *UL148*-null mutant virus used herein, which carries nonsense codons in *UL148*, phenocopied the enhanced epithelial cell tropism of the previously characterized *UL148* deletion mutant (20) (Fig. S2). Further, incorporation of the S-tag, which allows for S-affinity purification (S-AP) of gO during pulse-chase experiments, had no effect on viral replication kinetics even during low MOI infection (Fig. S1B). To visualize the global stability of gO, and to discriminate between immature and mature gO glycoforms, we subjected S-AP eluates to PNGase F and endoglycosidase H (endoH) digestion, respectively. After resolving SAP eluates by SDS-PAGE following pulse-chase and subjecting dried gels to autoradiography, we quantified the ~55 kD bands that corresponded to endoglycosidase-digested gO (Fig. 2).

**FIG. 2.**
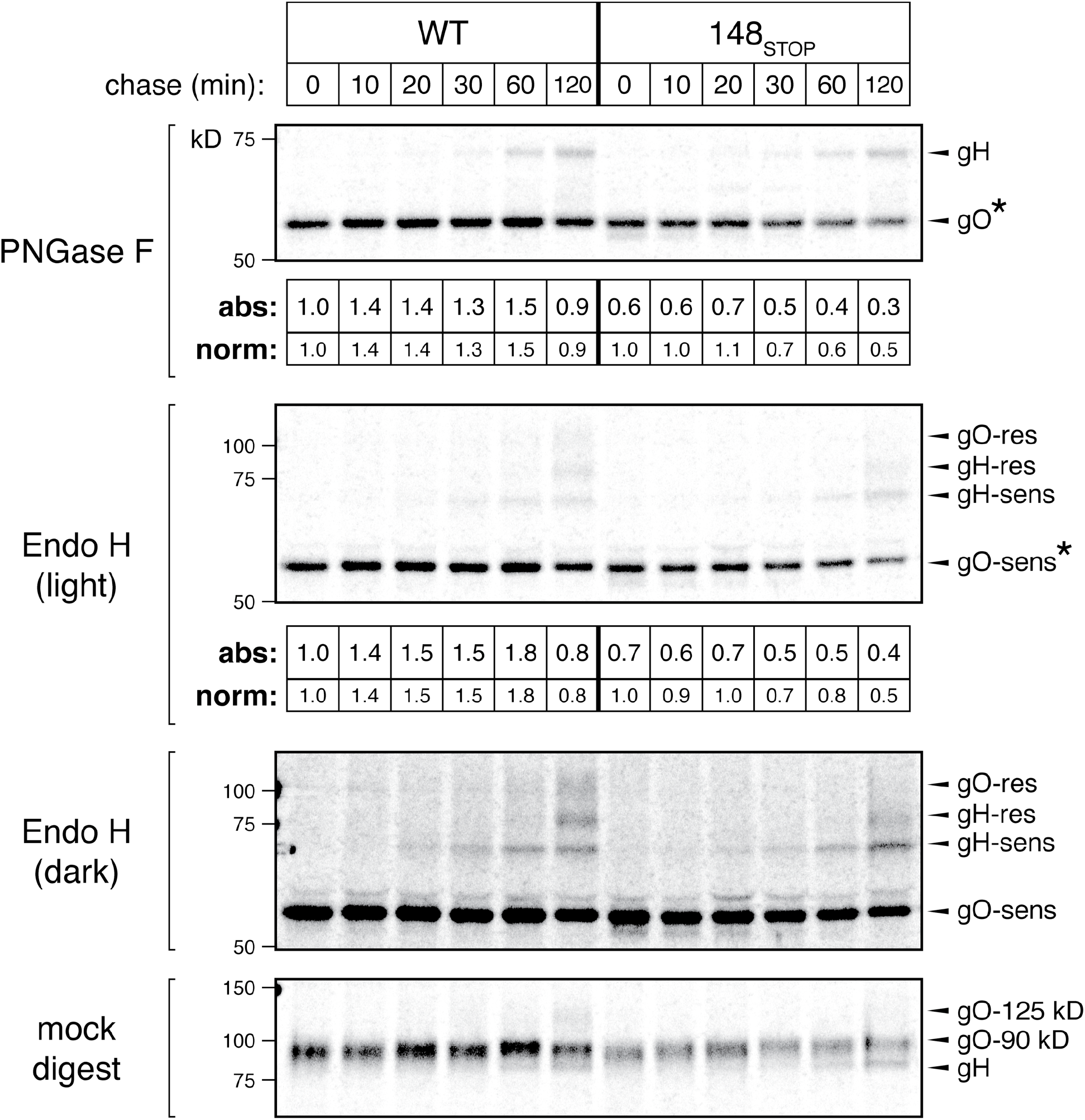
Pulse-chase analysis of gO during *UL148*-null HCMV infection. Fibroblasts (HFFTs) were infected at an MOI of 1 TCID_50_ per cell with TB40/E-derived recombinant TB_gO-S expressing S-tagged gO (WT) or its *UL148*-null derivative, TB_148_STOP__gO-S (148_STOP_). At 96 hpi, the cells were pulsed for 20 mins with 200 μCi/mL ^35^S-Met/Cys and chased for the indicated times before lysis. Equal TCA-precipitable CPMs of each sample were subjected to S-AP, followed by endoglycosidase digestion and SDS-PAGE. The dried gel was exposed to a phosphor screen to produce an autoradiograph. The densities of the indicated gO bands (*) were calculated and expressed either as absolute signal (“abs”) in relation to the WT/0 h band or as signal normalized (“norm”) across WT or *UL148*-null conditions relative to the respective 0 h band.

Notably, we found that gO decayed approximately 2-fold more rapidly during infection with the *UL148*-null virus, TB_148_STOP__gO-S (Fig. 2, “norm” signal in both endoH and PNGase F treatments). We also observed that infections with the wild-type (WT) comparator virus, TB_gO-S, exhibited higher absolute levels of labeled gO than TB_148_STOP_TOP_gO-S (Fig. 2, “abs” signal), even at the earliest chase time point (0 h). Weaker baseline signals for ERAD substrates have also been observed by others during conditions of more rapid decay, e.g. (30). We thus interpreted these results to indicate that gO undergoes accelerated ERAD in the absence of UL148.

### Knockdown of SEL1L leads to an increase in steady-state gO levels

We next sought to investigate whether gO is subjected to SEL1L-dependent ERAD during HCMV infection. To deplete cells of SEL1L, we transfected cells with siRNA pools targeting *SEL1L*, and then infected with either wild-type (TB_WT) or *UL148*-null mutant (TB_148_STOP_) HCMV strain TB40/E viruses. Depletion of SEL1L led to a robust increase in steady-state gO levels, while levels of other viral proteins such as pp150 and UL44 were found to be decreased or unaffected (Fig. 3). SEL1L knockdown likewise failed to stabilize any of the other three viral glycoproteins whose expression we monitored, gB, gH, and gL, and instead appeared to decrease their expression (Fig. 3). The observed decrease in levels of other viral gene products may due to the unfolded protein response (UPR), which can be activated during SEL1L depletion (31). Intriguingly, stabilization of gO in response to SEL1L siRNA treatment was more pronounced during WT virus infection than in the *UL148*-null setting.

**FIG. 3.**
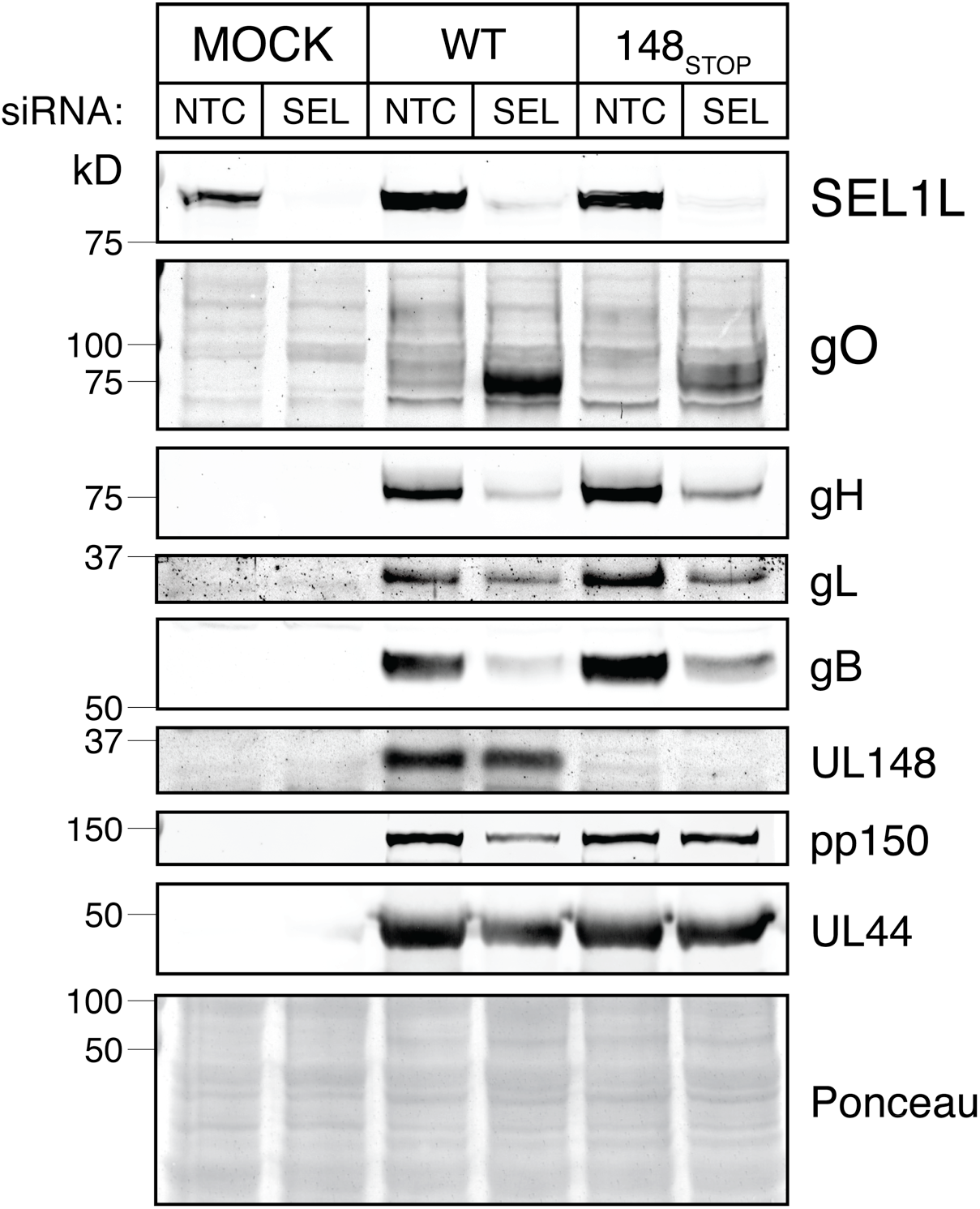
Depletion of SEL1L in HCMV-infected cells increases steady-state gO levels. (A) Fibroblasts (HFFTs) were reverse-transfected with siRNA targeting SEL1L (SEL) or non-targeting control (NTC). At 48 h post transfection, the cells were infected with HCMV strain TB40/E (WT) or its *UL148*-null derivative, TB_148_STOP_ (148_STOP_) at an MOI of 1 TCID_50_ per cell. HCMV glycoprotein levels at 96 hpi were analyzed by western blot.

Endoglycosidase H (EndoH) treatment, which removes N-linked glycans to cause ER glycoforms of gO to migrate as single species of M_r_ ~50 kD, confirmed that the major immunoreactive bands we detected during SEL1L depletion were indeed gO, and also indicated that SEL1L knockdown stabilized endoH-sensitive ER forms of gO, as would be consistent with the interpretation that gO is an ERAD substrate (Fig. S3A). Similar results were also seen when S-tag antibodies were used to detect SEL1L knockdown stabilization of gO in TB_gO-S infected cells (Fig. S3B). Arguing against the possibility that the effects of SEL1L depletion on gO levels might be a strain-specific phenomena, we found similar results for the HCMV laboratory strain AD169 [which lost *UL148* and several other viral genes during tissue culture adaptation (32)], and as was the case in WT vs. *UL148*-null TB40/E, we likewise observed amplified gO stabilization for an AD169 derivative restored for *UL148* (Fig. S4). Finally, siRNA knockdown of the SEL1L partner protein, the E3 ubiquitin ligase Hrd1, led to increased gO levels, as would be expected based on the effects of SEL1L knockdown (Fig. S5). We interpreted these results to strongly suggest that gO is a constitutive ERAD substrate during HCMV infection.

### Kifunensine treatment increases steady-state gO levels

We next tested whether pharmacological inhibition of ERAD would similarly stabilize gO. Kifunensine (KIF) is a potent inhibitor of ER mannosidase I (33), which demannosylates the N-linked glycans on misfolded glycoproteins to target them for recognition by ER lectins such as OS-9 and XTPB-3, which in turn deliver the misfolded glycoproteins to the SEL1L-Hrd1 complex for ERAD [(27), reviewed in (26)]. After treating HCMV-infected fibroblasts with KIF at 72 hpi and harvesting lysates at 96 hpi, we observed a robust, dose-dependent increase in the abundance of an intermediate-size gO glycoform that migrated at ~90 kD SDS-PAGE (Fig. 4, Fig. S6). Furthermore, as in the SEL1L and Hrd1 knockdown experiments, the presence of UL148 appeared to further enhance gO levels in both control and KIF-treated cells. In contrast to SEL1L depletion, however, we observed no significant decrease in levels of gH or gB after KIF treatment, suggesting that KIF may be less toxic to infected cells than siRNA depletion of SEL1L or Hrd1. These results further suggested to us that gO is a constitutive ERAD substrate during infection.

**FIG. 4.**
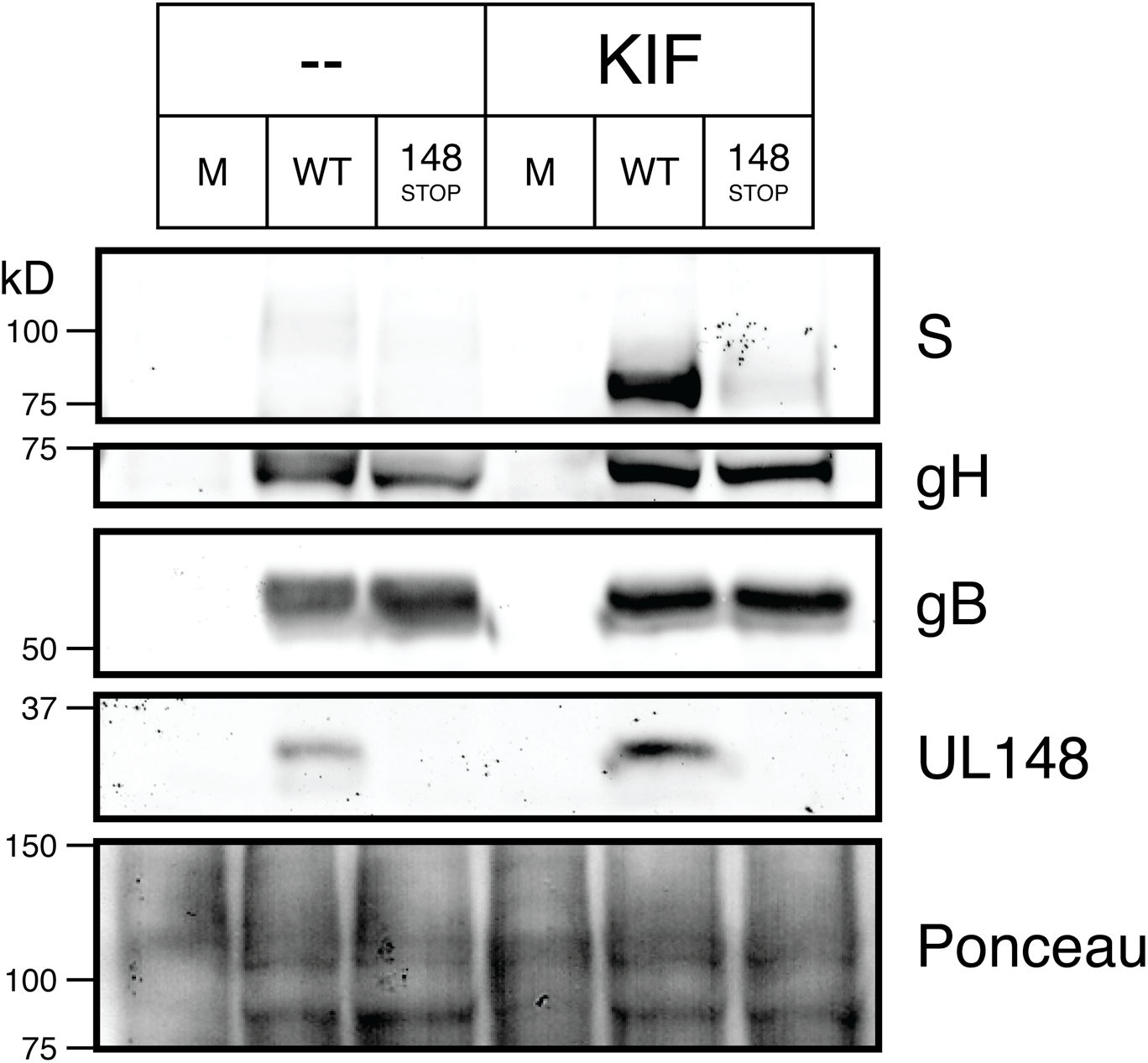
Treatment of HCMV-infected cells with an ERAD inhibitor increases steady-state gO levels. Fibroblasts (HFFTs) were infected at an MOI of 1 TCID_50_ per cell with TB_gO-S (WT), TB_148_STOP__gO-S (148_STOP_), or mock infection (M). Cells were treated with kifunensine (KIF) at 2.5 μM or carrier-alone (water) at 72 hpi. At 96 hpi, HCMV glycoprotein levels were analyzed by western blot.

### Pulse-chase analysis of gO during SEL1L and Hrd1 depletion

To evaluate how blockade of ERAD would affect gO decay in the presence versus the absence of UL148, we carried out pulse-chase experiments in fibroblasts that were siRNA depleted for SEL1L or Hrd1 and subsequently infected with WT or *UL148*-null viruses expressing S-tagged gO. To fully resolve gO from gH, we subjected S-AP eluates to PNGase F digestion. As expected, gO decayed more slowly during knockdown of either SEL1L or Hrd1 compared to non-targeting control (NTC) conditions, regardless of whether UL148 was present (Fig. 5A-B). Absolute levels of gO, however, were most markedly amplified—by up to 7-fold at the 4 h chase time point, when knockdown of either Hrd1, or to a lesser extent, SEL1L, occurred in presence of UL148 (Fig. 5B). The remarkable degree of gO stabilization during Hrd1 siRNA treatment made the roughly 2- to 3-fold effect of UL148 on gO stability, as seen in NTC treatments at 4 h post chase, appear modest by comparison (Fig. 5A-B). The observation that UL148 appeared to amplify stabilization of gO during ERAD blockade may suggest that UL148 acts upstream of Hrd1/SEL1L. Alternatively, this effect may simply reflect that substantial amounts of gO are still getting degraded by residual levels of SEL1L/Hrd1, despite efficient siRNA knockdown (Figs. 3, S4, S5), and that these residual levels of ERAD are readily squelched by UL148, which would be present at higher ratio to Hrd1 during the siRNA treatment. The latter possibility would be consistent with a functional relevance for the SEL1L interaction. Overall, these data corroborate the hypothesis that gO is targeted by ERAD during infection, and depending on the interpretation, might be taken to suggest that UL148 functions to inhibit ERAD of gO.

**FIG. 5.**
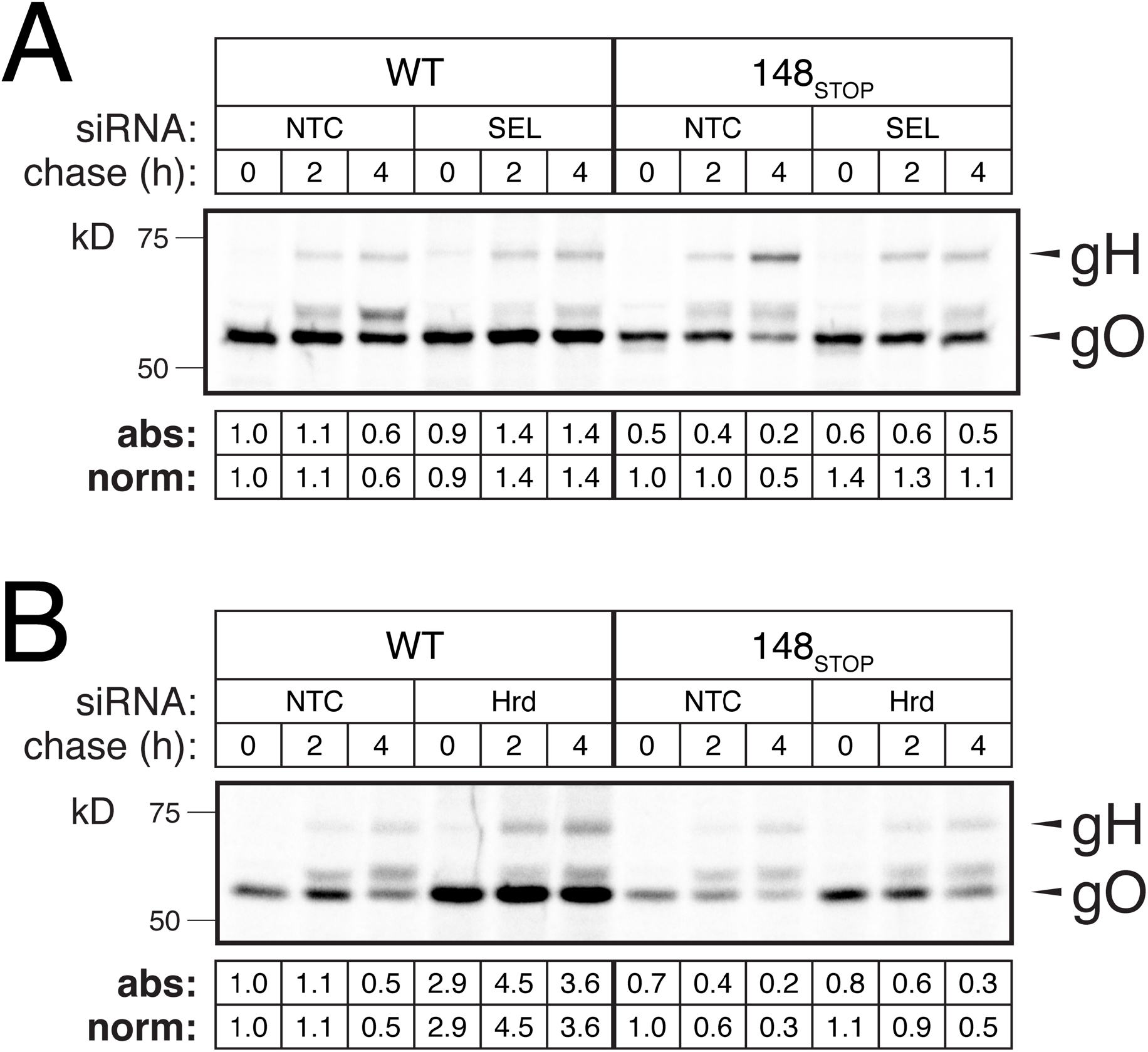
Depletion of SEL1L or Hrd1 in HCMV-infected cells stabilizes gO during WT and *UL148*-null infection. Fibroblasts (HFFTs) were reverse-transfected with siRNAs targeting (A) SEL1L or (B) Hrd1. At 6 hpt, the cells were infected at an MOI of 1 TCID_50_ per cell with TB_gO-S (WT) or TB_148_STOP__gO-S (148_STOP_). At 96 hpi, the cells were pulsed for 20 mins with 200 μCi/mL ^35^S-Met/Cys and chased for the indicated times before lysis. Equal TCA-precipitable CPMs of each sample were subjected to SAP, followed by PNGase F digestion to allow better resolution of gH and gO by SDS-PAGE. The dried gel was exposed to a phosphor screen to produce an autoradiograph. The densities of gO bands were calculated and expressed either as absolute signal (“abs”) in relation to the WT/NTC/0 h band or as normalized signal (“norm”) for each virus condition relative to the respective NTC/0 h band.

**FIG. 6.**
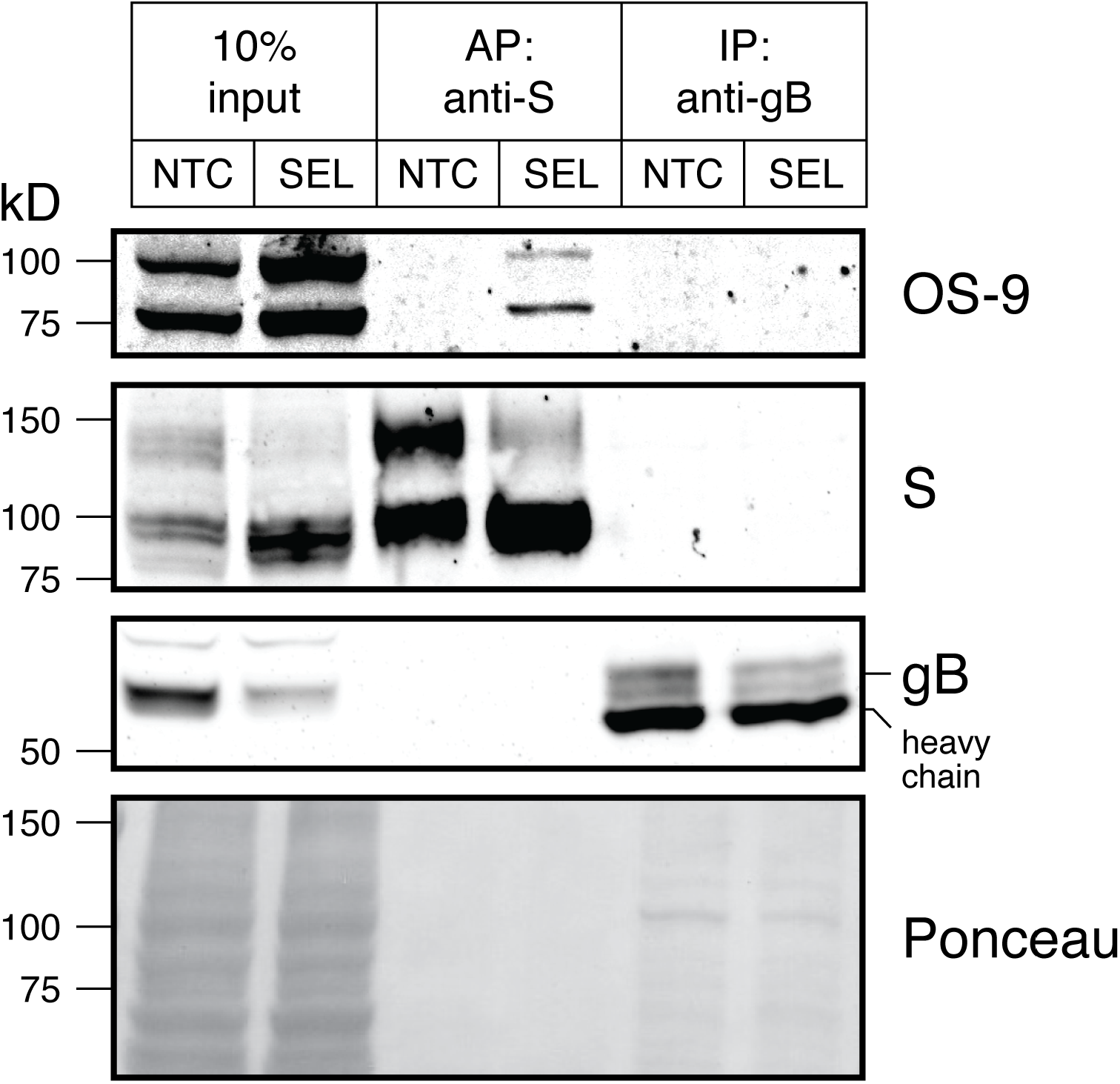
Depletion of SEL1L in HCMV-infected cells stabilizes a physical association between gO and ER lectin OS-9. Fibroblasts (HFFTs) were reverse-transfected with siRNAs targeting SEL1L (SEL) or NTC. 48 h later, the cells were infected with TB_gO-S at an MOI of 1 TCID_50_ per cell. At 96 hpi, cell lysates were subjected to S-AP or gB-IP and analyzed by western blot.

### Knockdown of SEL1L stabilizes an association between gO and the ER lectin OS-9

As a final complementary approach to verify whether gO behaves as an ERAD substrate during infection, we asked whether gO associates with an ER lectin involved in upstream events during ERAD. Depletion of SEL1L is known to stabilize what is otherwise a transient association of the ER lectin OS-9 with ERAD substrates (27). We thus tested whether gO associates with OS-9 during knockdown of SEL1L in the context of the HCMV infected cell. We failed to observe co-purification of OS-9 with S-tagged gO from infected cells that were treated with non-targeting control siRNA (NTC) (Fig. 5). In contrast, we repeatedly detected co-purification of OS-9 with gO-S from infected cells that were knocked down for SEL1L (e.g., Fig. 5). Furthermore, gB failed to co-IP OS-9 during either NTC or SEL1L knockdown, further arguing that the association that we detected between OS-9 and gO is specific. That we were able to detect co-purification of endogenous OS-9 with a putative ERAD substrate is notable. To our knowledge, recovery of detectable levels of endogenous OS-9 in this type of experiment has been observed only in one other study (34); most studies using this approach have relied upon OS-9 over-expression to enhance the sensitivity of the assay (27, 31, 35, 36). Overall, we interpreted this result to suggest that a physical association occurs between gO and OS-9 during HCMV infection, which further argues that gO is an ERAD substrate.

## DISCUSSION

Despite that this study was initiated to address the mechanisms underlying the influence of UL148 on trimer (gHgLgO) expression, one of the most striking findings is the unexpectedly large role for ERAD in regulating gO expression. Newly synthesized gO appears to be very unstable within the ER, much more so than other viral glycoproteins, as we did not find that pharmacologic or siRNA mediated inhibition of ERAD to stabilize any other viral glycoproteins. In fact, siRNA knockdown of either SEL1L or Hrd1 each led to decreased expression of gH, gL, and gB (Fig. 3) but amplified gO levels by up to 7-fold (Fig. 5B). Although the literature also suggests the existence of post-ER processes that regulate virion incorporation of gHgL complexes (37), because gO participates in alternative gHgL complexes that contribute to HCMV cell tropism, our findings underscore the large potential for ERAD to sculpt the glycoprotein composition of the virion envelope, and hence, to influence the infectivity of HCMV for different cell types.

### Why would gO be an ERAD substrate?

gO is a heavily glycosylated protein, with at least half of its ~100 kD molecular mass derived from N-linked glycans. Although the exact number varies between HCMV strains, its primary amino acid sequence contains approximately 20 N-x-(S/T) sequons that might serve as sites for N-linked glycosylation. Such a large degree of glycosylation may serve a number of purposes. For instance, gO also been reported to attenuate antibody-mediated neutralization of gH and gB (38). Hence, analogous to the “glycan shield” provided by gp120 in HIV (39), extensive glycosylation may contribute to the capacity of gO to protect the viral fusion machinery from neutralizing antibodies. However, given the roles that N-linked glycans play in glycoprotein quality control (26), the heavy degree of glycosylation may also predispose gO to be especially susceptible to ERAD targeting by ER mannosidases.

Additionally, inefficient folding may be an inherent property of the gO polypeptide. Previous pulse-chase studies have shown that gH forms a complex with gL within 30 min of its de novo synthesis, but that assembly of gHgLgO does not occur until approximately 2 h (40). This observation may imply that folding of gO occurs slowly and may be rate limiting for trimer assembly. Moreover, because gO must assemble into trimer in order to traffic further through the secretory pathway, the relatively slow kinetics of trimer assembly might compel a certain degree of constitutive degradation. Indeed, orphan subunits of multisubunit complexes, such as the T-cell receptor, are retained in the ER and destroyed via ERAD (41, 42). Competition with the pentamer-specific subunits UL128, UL130 and UL131 for assembly onto gHgL (9, 43–45) may further increase levels of unassembled gO that would targeted for disposal. Accordingly, upon repair of pentamer expression to HCMV strain AD169, Wang and Shenk showed decreased amounts of gO co-immunoprecipitated with gH [(9), Fig. 1C], and Zhou *et al*. found transcriptional suppression of the *UL128* locus to increase levels of gHgLgO in virions (21). It thus seems likely that both of these mechanisms, intrinsic instability and inefficient trimer assembly, may contribute to limiting trimer expression during HCMV infection.

Notably, we find that the homologs of gO from murine cytomegalovirus (MCMV) and rhesus cytomegalovirus also appear to be stabilized in infected cells during treatment with ERAD inhibitors or SEL1L knockdown (Zhang, H and Kamil, JP; unpublished results). Therefore, constitutive ERAD of gO homologs may prove to be a conserved feature among β-herpesviruses. Since the expression of alternative gHgL complexes is also shared across the β-herpesvirus subfamily, the intrinsic instability of gO and its homologs could provide a basis for mechanisms to modulate the composition of alternative gHgL complexes in a cell-type specific manner.

### How does UL148 influence ERAD of gHgLgO?

Our results suggest that gO is targeted by ERAD during infection, and that UL148 limits its degradation, with implications for HCMV cell tropism that are summarized in our model (Fig. 7). UL148 could accomplish its effects on gO by at least two distinct but related mechanisms: (i) by promoting the folding and/or assembly of gO into mature trimer complexes, thereby preventing gO from being shunted to ERAD, or (ii) by dampening the ERAD pathway generally, perhaps through an interaction with SEL1L. The first mechanism would be consistent with our previous data suggesting that UL148 interacts with immature gHgL complexes during their transit through the ER (20). Although we cannot yet exclude the second possibility, determining whether UL148 stabilizes classical ERAD substrates is a foremost experimental priority. Moreover, the two mechanisms are not mutually exclusive. For instance, UL148 may bind to the gHgL dimer and inhibit the ERAD machinery to allow time for gO to assemble onto the complex. Interestingly, we consistently observed a UL148-dependent increase in the absolute levels of gO, whether or not ERAD was disrupted, which might be taken to favor the possibility that UL148 enhances gO levels via mechanisms upstream of ERAD. On the other hand, the observation of enhanced gO stabilization when UL148 is present during ERAD blockade may simply reflect an influence of UL148 against the residual levels of ERAD that remain after siRNA- or pharmacologic disruption.

**FIG. 7.**
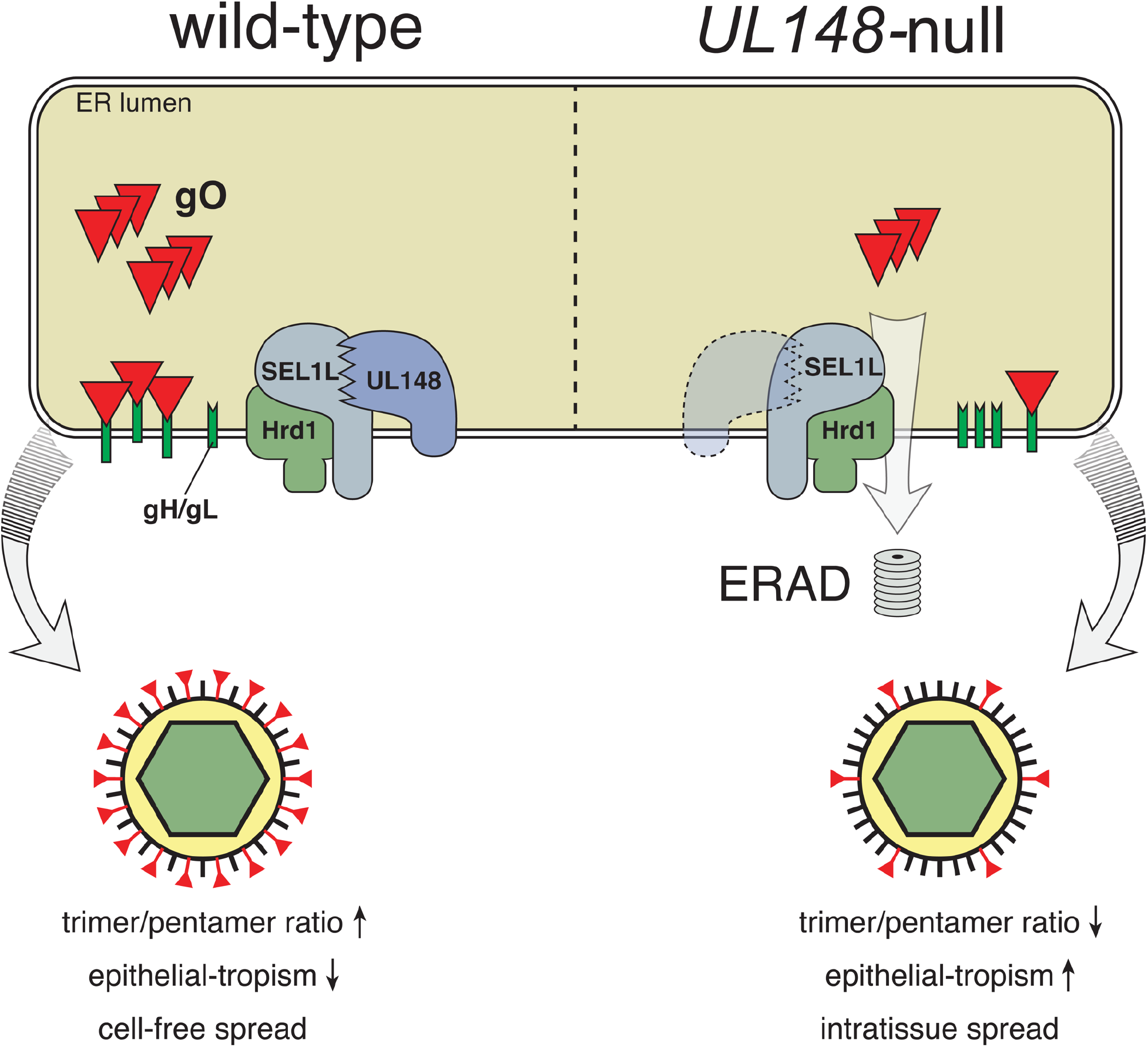
Model. HCMV gO, either through unfavorable folding or inefficient assembly onto gH/gL, is targeted for destruction by ERAD, which is mediated by SEL1L/Hrd1. UL148, perhaps through a functional interaction with SEL1L, slows the degradation of gO.

### Implications of gO instability and trimer regulation by UL148

Tissue culture models have consistently demonstrated the importance of gHgLgO for infection of several cell types (12, 13, 22). However, *in vivo* studies using MCMV suggest that while the analogous complex of m74 with gHgL is required for initial infection of tissues by cell-free virions, it is dispensable for subsequent intra-tissue spread (17). By analogy, it seems plausible that *in vivo* most HCMV-infected cells constitutively degrade gO to promote “immunologically covert” cell-associated modes of virus spread that are driven by the pentameric gHgL complex (11), thereby mimicking the *UL148*-null phenotype shown in our model (Fig. 7). In cell types relevant for horizontal spread, however, UL148 expression or activity may be enhanced to promote higher levels of gO, as would be consistent with previous reports suggesting that the type of cell producing virions influences the processing of virion components and virion assembly to yield distinct virus populations (18, 19). In support of this notion, the repaired HCMV strain Merlin (46)—which is thought to recapitulate features of HCMV that are rapidly lost during adaptation to growth in cultured cells, and which expresses trimer very poorly, is found to express UL148 at much lower levels than a HCMV clinical strain known to express high levels of trimer (47). Given that HCMV is highly cell-associated *in vivo* (2), it is intriguing to consider that UL148 abundance might be regulated so that trimer-rich virions are produced only in settings such as glandular or ductal epithelial cells where production of cell-free virions would be necessary for horizontal spread.

## MATERIALS AND METHODS

### Cells

Human foreskin fibroblasts (HFF) were immortalized by transducing primary HFFs (ATCC #SCRC-1041) with lentivirus encoding human telomerase (hTERT) to yield HFFT cells (see below). 293T cells were purchased from Genhunter Corp. (Nashville, TN). ARPE-19 retinal pigment epithelial cells were purchased from ATCC (CRL-2302). All cells were cultured in Dulbecco’s Modified Eagle’s Medium (DMEM, Corning) supplemented with 25 μg/mL gentamicin (Invitrogen), 10 μg/mL ciprofloxacin (Genhunter), and either 5% fetal bovine serum (FBS, Sigma #F2442) or 5% newborn calf serum (NCS, Sigma #N4637).

### Viruses

Virus was reconstituted by electroporation of HCMV bacterial artificial chromosomes (BACs) into HFFTs, as described previously (20, 48). HCMV strain AD169rv repaired for *UL131* (ADr131) (20) and recombinants derived from it were amplified at low MOI on ARPE-19 cells until 100% cytopathic effect (CPE) was observed. Virus-containing culture supernatants were then subjected to centrifugation (1000*g*) for 10 minutes to pellet any cellular debris. The supernatant was then ultracentrifuged (85,000*g*, 1 h, 4°C) through a 20% sorbitol cushion, and the resulting virus pellet was resuspended in DMEM containing 20% NCS. For TB40/E and related recombinants, virus was grown on HFFTs until 100% CPE was observed. Cell-associated virus was then released by Dounce-homogenization of pelleted infected cells, clarified of cell debris by centrifugation (1000*g*, 10 min), and combined with the cell-free medium before ultracentrifugation and resuspension as above.

### Virus titration

Infectivity of virus stocks and samples were determined by the tissue culture infectious dose 50% (TCID_50_) assay. Briefly, serial dilutions of virus were used to infect multiple wells of a 96-well plate. After 9 days, wells were scored as positive or negative for CPE, and TCID_50_ values were calculated according to the Spearman-Kärber method (49, 50).

### Virus growth kinetics

For viral yield on fibroblasts (HFFT) vs ARPE-19 epithelial cells (Fig. S1), cells were seeded in a 24-well plate at 6.5 × 10^4^ cells per well. For multi-cycle growth kinetics of TB_gO-S (Fig. S3), HFFT were seeded at 1.5 × 10^5^ cells per well. Wells were then infected in duplicate at the indicated MOIs in in 0.3 mL per well, and back-titration of the inocula was initiated in parallel. Inocula were removed after 16 hr, and the cells were washed three times with 1 mL of Dulbecco’s phosphate-buffered saline (DPBS, Lonza Biowhittaker, Inc.). Cell-free supernatants were collected at indicated times post-infection and stored at −80°C until analysis. Infectivity titers (TCID_50_) were determined in parallel.

### Antibodies

Mouse monoclonal antibodies (mAbs) specific for gH (AP86) (51), gB (27-156) (52), and pp150 (53) were generous gifts from William J. Britt (University of Alabama, Birmingham). Additional gB and UL44 mouse mAbs were purchased from Virusys (#P1201 and #P1202-2). Rabbit antibodies were used to detect the following proteins: UL148 (20), SEL1L (Sigma #S3699), OS-9 (Cell Signaling Tech #12497), HA epitope tag: YPYDVPDYA (Bethyl #A190-108A), S-peptide: KETAAAKFERQHMDS (Bethyl #A190-135A). Additional rabbit sera immunoreactive to gL, and to gO variants from HCMV strains TB40/E and AD169 were generous gifts from Brent J. Ryckman (University of Montana, Missoula, MT) (21).

### Construction of recombinant viruses

Synthetic dsDNAs (gBlocks) and oligonucleotides were purchased from Integrated DNA Technologies (Coralville, IA); full sequence details are provided in Table S2. New recombinant HCMVs were constructed in the context of the infectious bacterial artificial chromosome (BAC) clones of HCMV strains TB40/E, TB40-BAC4 (54), and AD169, AD169rv (55), as previously described (20, 48, 56), with full details provided in Supplemental Text S1. Briefly, two-step Red *“en passant’* recombination (57, 58) was used in conjunction with GS1783 *E. coli* (a gift of Greg Smith, Northwestern University) harboring BAC-cloned HCMV genomes. Recombinant BACs were confirmed by Sanger-sequencing (Genewiz, Inc.) of modified loci (not shown), and by BamHI and EcoRI restriction analysis (not shown) to exclude spurious rearrangements.

*TB40/E expressing UL148^HA^ or UL16^HA^*. TB_148^HA^ was described previously (20). Text S1 describes construction of recombinant TB_UL16^HA^.

*UL148-null recombinant TB_148_STOP_*. TB_148_STOP_ was constructed by replacement of the UL148 coding sequence (CDS) with a synthetic version in which all in-frame methionine codons were replaced with nonsense codons.

*TB40/E and TB_148_STOP_ expressing S-tagged gO*. To add a bovine pancreatic ribonuclease A S-peptide tag (S-tag) (59) (KETAAAKFERQHMDS) to the *UL74* (gO) ORF without disrupting the overlapping gene, *UL73 (gN)*, a codon-optimized sequence encoding the following elements was inserted between nucleotides 139966 and 139967 (numbering per Genbank Accession #EF2999921.1): (i) the 30 C-terminal amino acids of gO, (ii) triple (Gly-Gly-Gly-Ser) linker, and (iii) the S-tag, followed by a tandem pair of stop codons.

*ADr131 expressing UL148^HA^*. Repair of *UL131* in strain AD169rv (ADr131) was described previously (20). ADr131_148 was constructed by replacing the vestigial *UL148* remnant with full-length *UL148* from HCMV strain TB40/E.

### Plasmids

The construction of new plasmids for this study is described in Text S1.

### hTERT lentivirus production and transduction

hTERT lentivirus was isolated and used to transduce HFF according to a modified version of the Addgene pLKO.1 protocol, as described previously (48). See Text S1 for details.

### siRNA treatments

siGenome SMARTpool siRNAs targeting SEL1L, Hrd1, and non-targeting control (NTC) siRNA were purchased from Dharmacon. Details are provided in Table S3. siRNAs were reverse-transfected into cells using Lipofectamine RNAiMAX reagent (Thermo Fisher) as per the manufacturer’s instructions. Briefly, two mixes were prepared separately: Mix #1 was prepared by adding 30 pmol of siRNA to 150 μL non-supplemented OptiMEM medium (Thermo Fisher) and gently mixing. Mix #2 was prepared by adding 9 μL of RNAiMAX reagent to 150 μL non-supplemented OptiMEM. Mix #1 and #2 were then combined and immediately transferred to an empty well of a 6- well plate and incubated at room temperature for 5 min. Approximately 1.0 million cells were then added to the well in 2.2 mL of DMEM containing 5% NCS (10 nM siRNA final).

### Endoglycosidase H analysis

EndoH digestion of 20 μg total protein was performed according to the manufacturer’s instructions (New England Biolabs).

### Pulse-chase

4 × 10^6^ HFFTs were infected at an MOI of 1 TCID_50_ per cell with TB_gO-S and TB_148_STOP__gO-S. In experiments involving knockdown of SEL1L or Hrd1, the cells were first reverse-transfected with siRNAs targeting SEL1L, Hrd1, or NTC, as described above; at 8 hours post-transfection (hpt), the cells were washed with DPBS and infected as above. At 96 hpi, the cells were washed twice with 10 mL DPBS and incubated for 30 mins in 5 mL of Met/Cys starving medium [DMEM lacking Met, Cys, and Glu (Gibco #21013024) plus 2 mM glutamate and 5% dialyzed FBS (Sigma #F0392)]. Cells were then pulsed for 20 mins with 2 mL of starving medium plus 200 μCi/mL ^35^S-Met/Cys (PerkinElmer #NEG772). To quench radiolabeling, 6 mL of chase medium [DMEM (Corning #10013CV) plus 2 mM Met, 2 mM Cys, and 5% NCS] was added directly to each dish. At the indicated time-points, the chase medium was replaced with ice-cold PBS. Cells were scraped, pelleted (4000*g* for 1 min), and lysed in RIPA buffer containing 25 mM HEPES (pH 7.5), 400 mM NaCl, 0.1% SDS, 0.5% sodium deoxycholate, 1% NP-40, 1% bovine serum albumin (BSA), and 1X protease inhibitor cocktail (PIC, CST #5871). Lysis continued for 1 h (4°C with rotation), and lysates were clarified of insoluble material by centrifugation (20,000*g*, 30 mins) and discarding of the pellet.

To quantify radiolabel incorporation into protein, 10 μL of lysate was added to 1 mL of 10% trichloroacetic acid (TCA) in triplicate, vortexed, and incubated on ice for 30 mins. Lysate/TCA solution was pipetted onto a glass disk (Whatman #1821025), and the disk was washed with 10 mL ice-cold 10% TCA and 10 mL ice-cold 100% ethanol. Disks were air-dried for 30 mins and shaken overnight in 5 mL scintillation fluid (Fisher #SX12-4). Counts per minute (CPMs) were measured on a scintillation counter (Beckman LS 6500) and averaged across triplicates.

#### S-protein affinity purification of radiolabeled gO

Equal CPMs of lysate were combined with 30 μL of S-protein agarose bead slurry (EMD Millipore) and RIPA buffer to a final volume of 400 μL. S-AP mixes were rotated overnight at 4°C. Beads were pelleted (1500*g*, 5 mins) and washed twice with 400 μL RIPA buffer containing 1% BSA, then washed twice with 400 μL RIPA buffer without BSA.

#### Endoglycosidase-digestion of S-AP eluates and autoradiography

To more effectively resolve co-purified gO and gH during SDS-PAGE, S-AP eluates were first digested with either peptide:N-glycosidase F (PNGase F, NEB) or endoglycosidase H (Endo H, NEB) as follows. After the final wash of S-AP above, the beads were boiled (85°C, 5 min) in 1X glycoprotein denaturing buffer, chilled on ice, then incubated with PNGase F or Endo H digestion mix (final volume 21 μL, 37°C, 1 h) according to the manufacturer’s instructions. 10 μL of 4X Laemmli buffer containing 20% β-mercaptoethanol (BME) was added to each sample, followed by boiling (85°C, 5 mins). Samples were resolved by sodium dodecyl sulfate-polyacrylamide gel electrophoresis (SDS-PAGE), as described previously (48). Gels were dried using a Bio-Rad Model 583 Gel Dryer, and radioactive signal was captured on a storage phosphor screen for a minimum of 72 h prior to data acquisition on Bio-Rad Molecular Imager FX. Band intensities were quantified using Quantity One 1-D Analysis Software (Bio-Rad). In brief, volume rectangles were drawn closely around the 55 kD gO band to generate a report of adjusted volume (CNT*mm^2^) values after local background subtraction. These values were normalized and reported as “gO signal”.

### Mass spectrometry

2 × 10^7^ fibroblasts (HFF) were infected at an MOI of 1 TCID_50_ per cell with TB_148^HA^. Cells were lysed at 120 hpi in modified RIPA lysis buffer [50 mM HEPES (pH 7.5), 1% TritonX-100, 400 mM NaCl, 0.5% sodium deoxycholate, 10% glycerol, and 1X PIC]. Anti-HA magnetic beads (Pierce #8837) were incubated (rotation overnight, 4°C) with lysates plus 100 μg/mL BSA, washed three times with lysis buffer, and eluted by boiling in Laemmli buffer (65°C, 5 min). Eluate was resolved by SDS-PAGE, and the gel was silver-stained according to the manufacturer’s protocol (ThermoFisher #24600). Silver-stained bands were excised from SDS-PAGE gels and sent to the Taplin Biological Mass Spectrometry Facility at Harvard Medical School (Boston, MA). Processing of gel slices for trypsin digestion, and resolution of peptides by nano-scale capillary reverse-phase HPLC are described in SI Methods. Eluted peptides were subjected to electrospray ionization and entered an LTQ Orbitrap Velos Pro ion-trap mass spectrometer (Thermo Fisher). Peptides were fragmented, and specific fragment ions were detected to generate a tandem pass spectrum for each peptide. Each spectrum was matched to a fragmentation pattern database by the program Sequest (Thermo Fisher) to determine the sequence of each peptide and hence protein identity (60). All databases included forward and reverse versions of peptide sequences, and data was filtered on a 1-2% peptide false discovery rate.

### Immunoprecipitation from transfected cells

5 × 10^5^ 293T cells were transfected with 2.5 μg of pEF1α V5 C vector (Invitrogen) encoding codon-optimized *US11^HA^, UL16^HA^*, or *UL148^HA^* using Mirus TransIT-2020 (#MIR 5404) according to the manufacturer’s instructions. At 48 hpt, cells were lysed in 300 μL of lysis buffer [50 mM HEPES (pH 7.5), 400 mM NaCl, 0.5% sodium deoxycholate, and 1X PIC]. 200 μL of lysate was rotated overnight (4°C) with 25 μL of anti-HA magnetic bead slurry (Pierce #8837). Beads were washed three times with lysis buffer and eluted by heating (50°C, 10 min) in 2X Laemmli buffer containing 5% (v/v) BME. Eluates and whole cell lysates were analyzed by western blot.

### OS-9 co-purification assay

1 × 10^6^ HFFT were reverse-transfected with siRNAs targeting SEL1L or NTC, as above. At 48 hpt, cells were infected with TB_gO-S virus at an MOI of 1 TCID_50_ per cell. At 96 hpi, cells were lysed in 50 mM HEPES (pH 7.5), 1% TritonX-100, 400 mM NaCl. Equal μg of protein (determined by Pierce BCA assay) were subjected to S-AP [30 μL of S-protein agarose slurry] or glycoprotein B (gB)-IP [30 μL Protein G magnetic bead slurry (EMD Millipore) plus anti-gB mAb 27-156 (52)]. After overnight rotation (4°C), beads were washed three times with lysis buffer and eluted in 1X Laemmli buffer plus 5% BME. Eluates and whole cell lysates were analyzed by western blot.

## ACKNOWLEDGMENTS

This project was supported by NIH Grants R01-AI116851 and P30GM110703. Its contents are solely the responsibility of the authors and do not necessarily represent the official views of the funding agencies. C.C.N. was supported by a fellowship from the Center of Cardiovascular Diseases and Sciences at LSU Health Sciences Center - Shreveport. We thank William Britt, Brent Ryckman, Greg Smith, Christian Sinzger, and Ulrich Koszinowski for generously sharing reagents.

## Author contributions are as follows

C.N. performed the majority of experiments. C.C.N. and J.P.K. designed experiments, interpreted results, and wrote the manuscript. J.P.K. designed HCMV recombinants. M.N.S. constructed recombinant TB_148_STOP_, performed UL148/SEL1L and gO/OS-9 co-purification studies, and prepared samples for MS analysis. H.Z. constructed recombinant viruses TB_gO-S and TB_148_STOP__gO-S and assisted with replication of virus yield experiments.

---

**FIG. S1.**
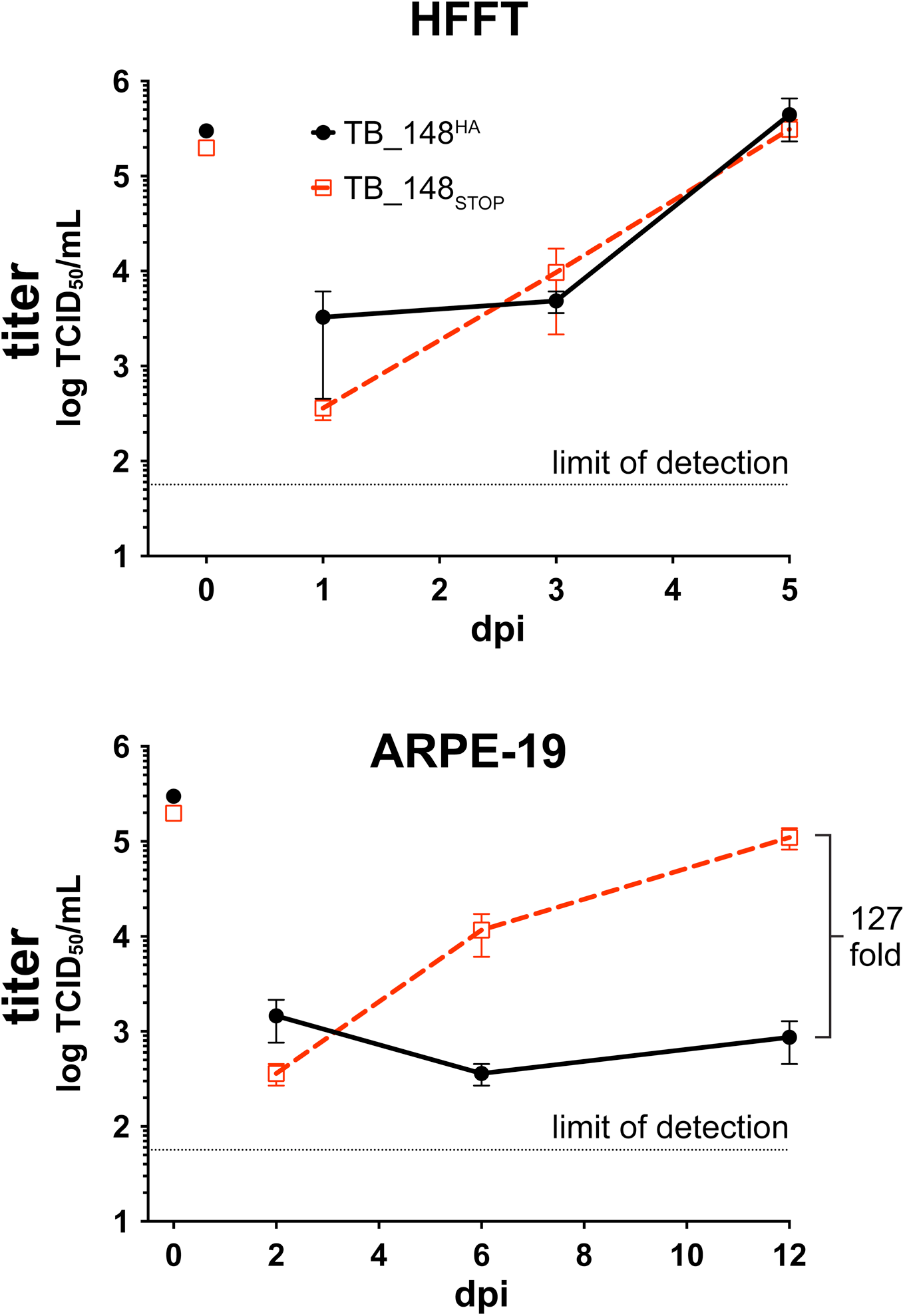
Construction and characterization of TB_gO-S. (A) Strategy for adding a C-terminal S-tag to the *UL74* (gO) ORF with minimal disruption of essential ORF *UL73*. (B) Multi-cycle growth kinetics of TB_gO-S. Fibroblasts (HFFT) were infected with TB40/E (TB) or TB_gO-S at an MOI of 0.005 TCID_50_ per cell. Cell supernatants were collected at the indicated dpi and titered in parallel by TCID_50_ assay. Back-titration of the original infectious inocula is depicted at the 0 dpi time-point. Error bars depict SEM calculated from duplicate infected wells. Dataset depicts one of two independent experiments.

**FIG. S2.**
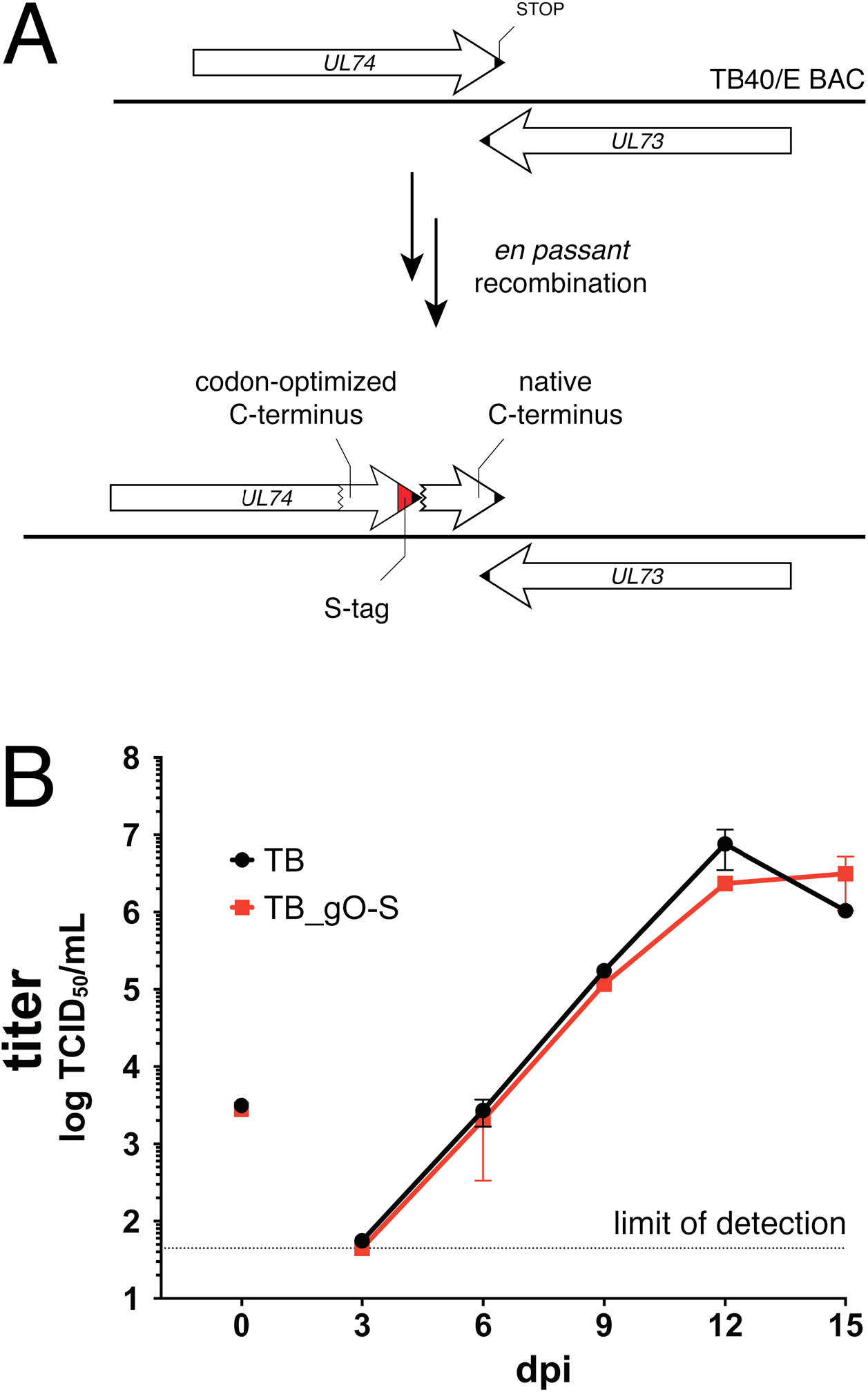
Growth kinetics of TB_148^HA^ and TB_148_STOP_ on HFFT and ARPE-19 cells. Fibroblasts (HFFT) and ARPE-19 cells were infected with TB_148^HA^ and TB_148_STOP_ at an MOI of 1 TCID50 per cell according to virus stock titers calculated on HFFTs. Cell-free supernatants were collected at the indicated dpi and titered in parallel by TCID_50_ assay on HFFTs. Back-titration of the original infectious inocula is depicted at the 0 dpi time-point. The dataset shown is one of two independent experiments. Error bars depict SEM calculated from duplicate infected wells.

**FIG. S3.**
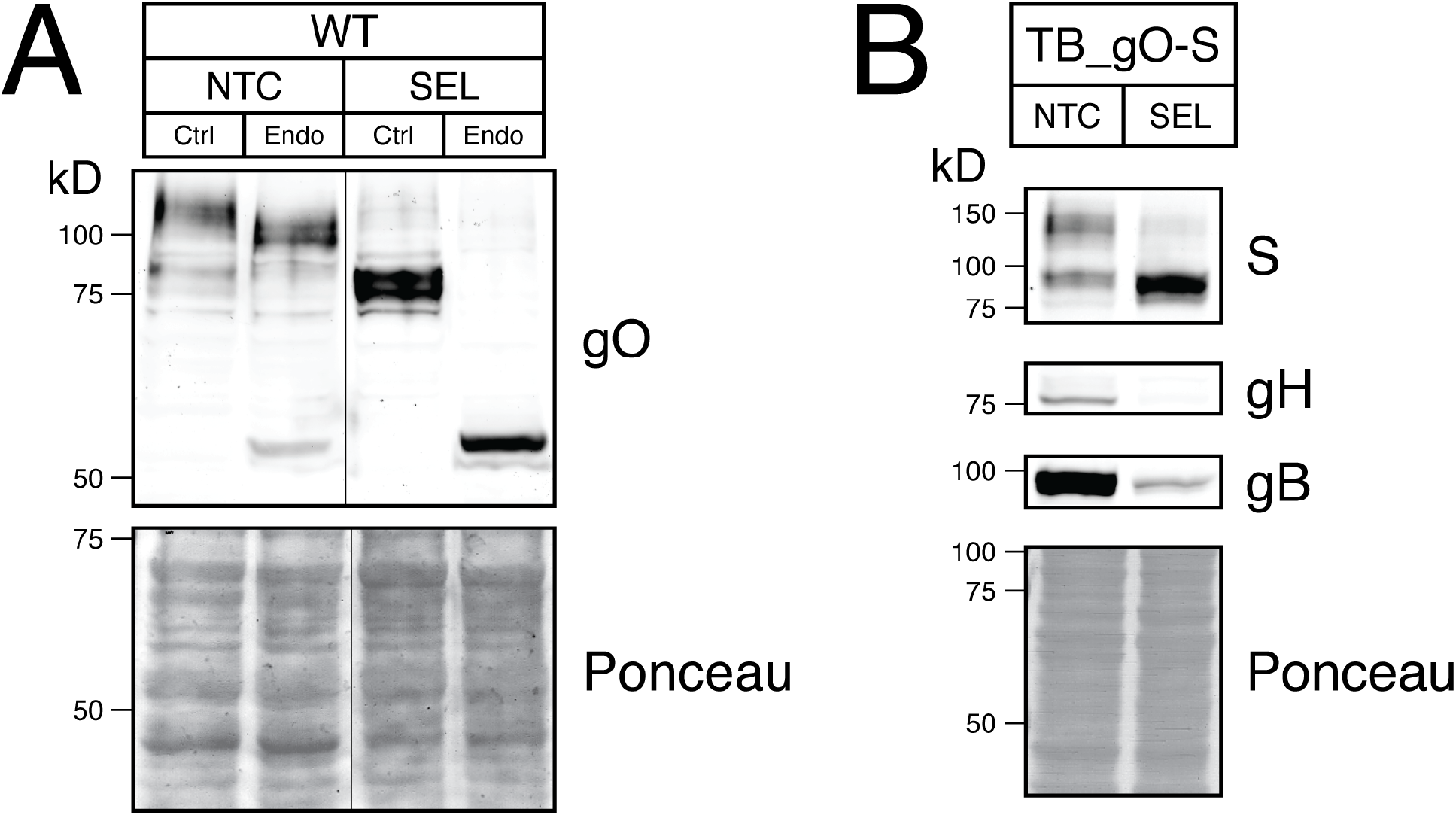
EndoH analysis and detection of gO levels during SEL1L knockdown. (A) Lysates from TB_WT-infected, siRNA-transfected cells (Fig. 3) were treated with EndoH or mock-digest. Extraneous lanes from this blot have been cropped from the image. (B) S western blot analysis of HCMV-infected cells depleted of SEL1L. Fibroblasts (HFFT) were reverse-transfected and infected as in Fig. 3 with TB_gO-S. Lysates were harvested at 96 hpi and analyzed by western blot.

**FIG. S4.**
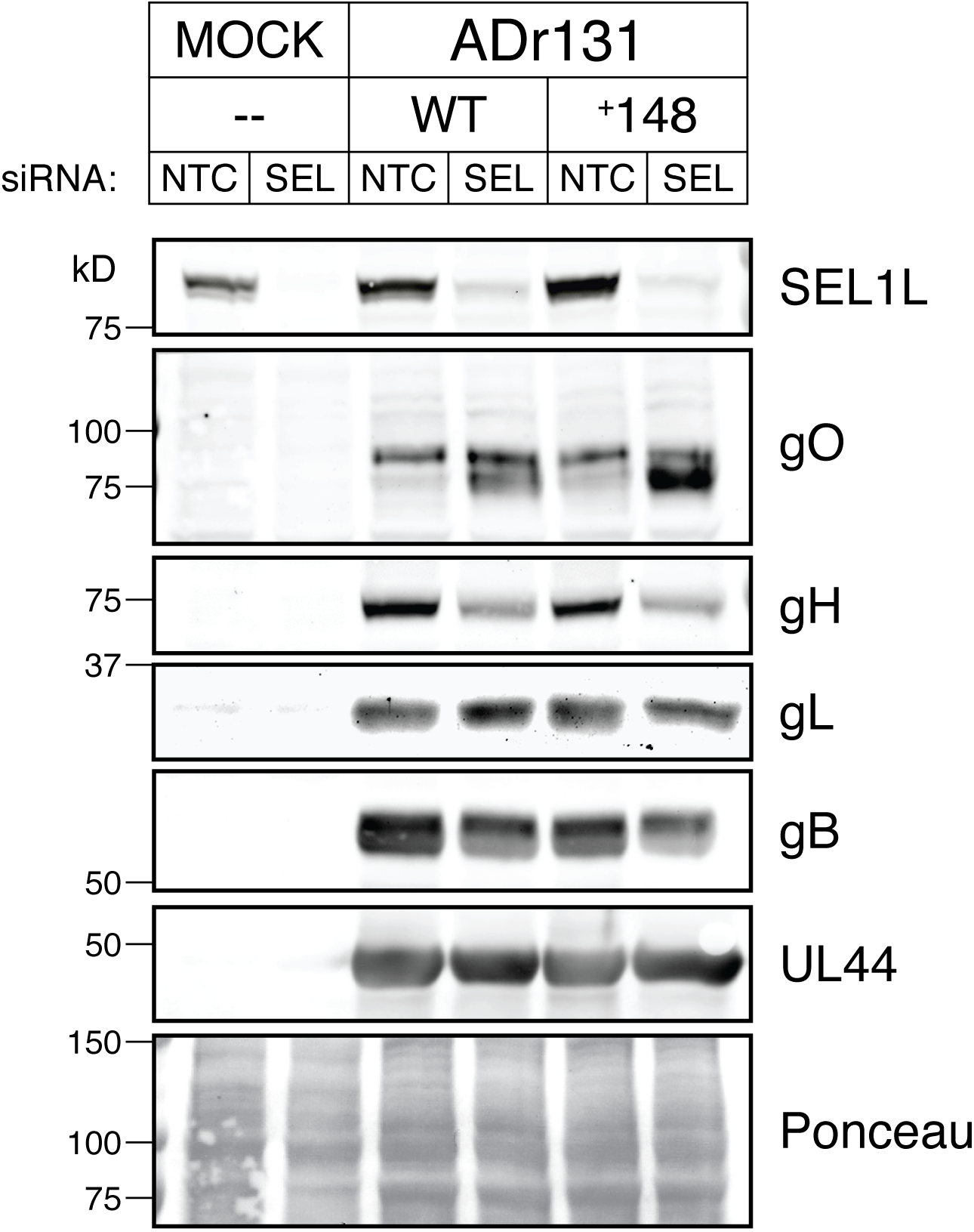
Depletion of SEL1L in HCMV strain AD169-infected cells recapitulates effects on HCMV glycoproteins. Fibroblasts (HFFT) were reverse-transfected and infected as in Fig. 3 with recombinant derivatives of HCMV strain AD169: ADr131 (WT) or ADr131_148 (^+^148). Lysates were harvested at 96 hpi and analyzed by western blot.

**FIG. S5.**
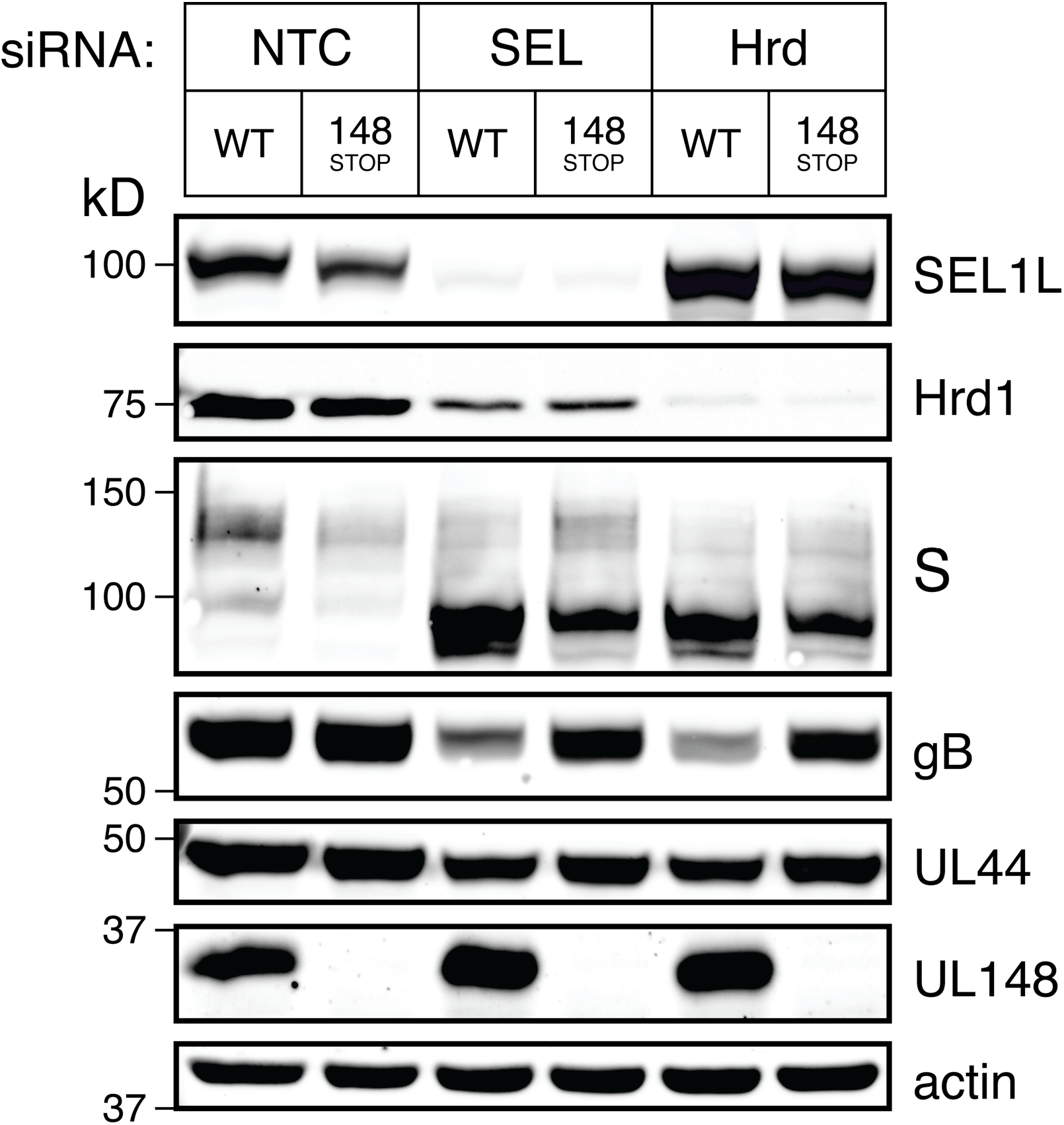
Hrd1 depletion in HCMV-infected cells increases gO levels. Fibroblasts (HFFT) were reverse-transfected siRNAs against NTC, SEL1L (SEL), or Hrd1 (Hrd) and infected with TB_gO-S (WT) or TB_148_STOP__gO-S (148_STOP_), as in Fig. 3. Lysates were harvested at 96 hpi and analyzed by western blot.

**FIG. S6.**
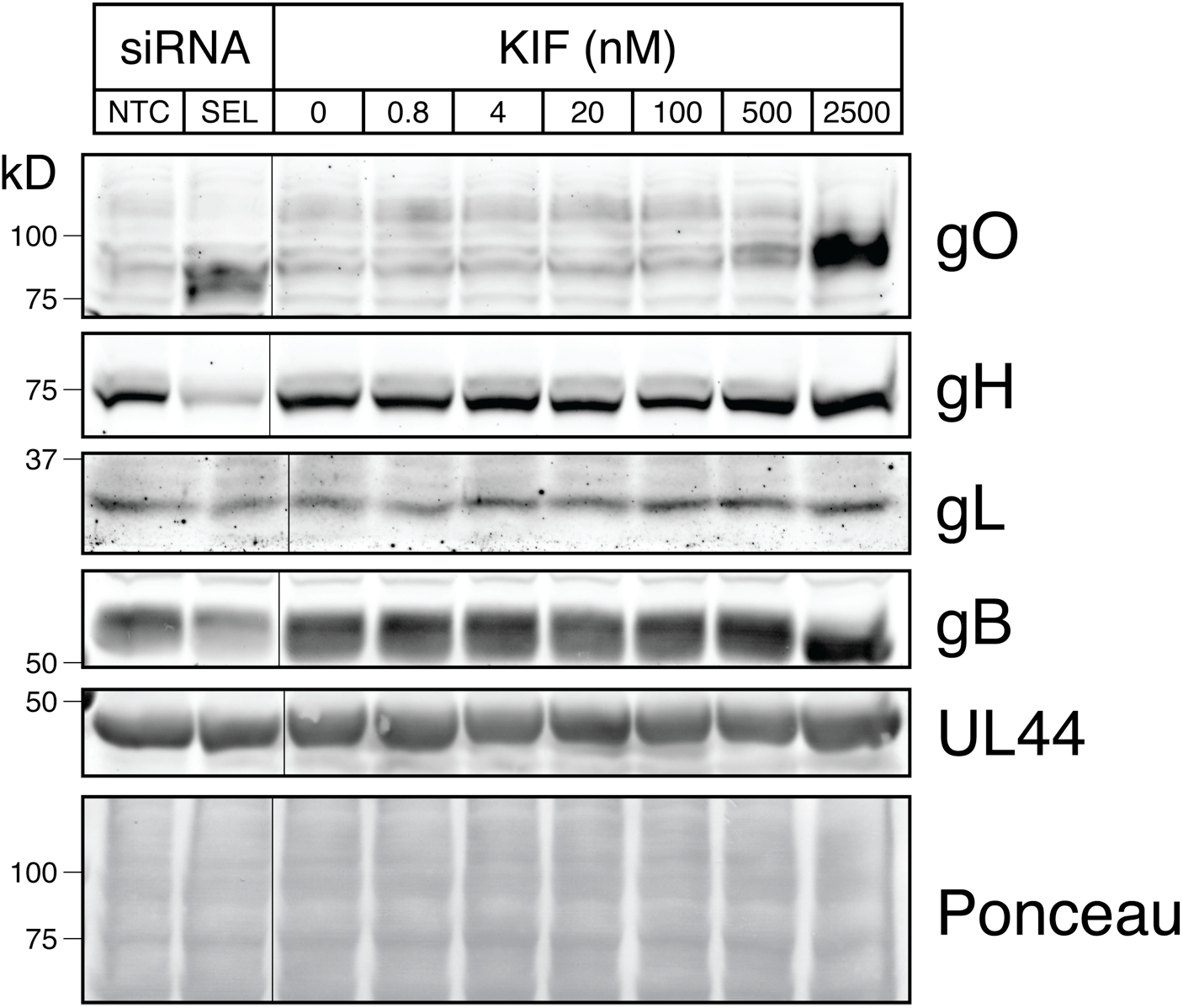
Dose-responsive effect of kifunensine on HCMV gO levels. Fibroblasts (HFFT) were infected with HCMV strain TB40/E at an MOI of 1 TCID_50_ per cell. At 72 hpi, fresh medium containing the indicated concentrations of KIF were added to the cells. Lysates were harvested at 96 hpi and analyzed by western blot. gO was visualized by antiserum against TB40/E gO. As a positive control for steady-state gO increase, cells were reverse-transfected with anti-SEL1L siRNAs as in Fig. 3, were infected, and analyzed in parallel. Extraneous lanes have been cropped out of the image. Concentrations above 2500 nM did not further increase gO levels (data not shown).

**TABLE S1.** Mapping of peptide hits from UL148HA-IP eluates to human gene products. Dataset depicts the unique and total hits mapping to the protein products of the human genes in the left column (HGNC Database)(6). Candidate genes were included in the list only if they appeared in both TB_148^HA^-IP replicates and did not appear in the TB_UL16^HA^-IP negative control.

| Gene Symbol | Replicate 1 |  | Replicate 2 |  |
| --- | --- | --- | --- | --- |
|  | Unique | Total | Unique | Total |
| ATP2A2 | 2 | 2 | 3 | 3 |
| CALU | 4 | 4 | 2 | 2 |
| CAND1 | 3 | 3 | 7 | 7 |
| CD58 | 1 | 1 | 1 | 1 |
| CLPTM1 | 1 | 1 | 3 | 3 |
| COPB1 | 1 | 1 | 2 | 2 |
| COPG1 | 5 | 5 | 5 | 6 |
| CTNND1 | 4 | 4 | 5 | 5 |
| DDOST | 5 | 5 | 4 | 4 |
| DHCR7 | 5 | 6 | 4 | 4 |
| DKK3 | 1 | 1 | 2 | 2 |
| DNAJB11 | 3 | 3 | 2 | 2 |
| DNAJC3 | 7 | 7 | 7 | 8 |
| ERLEC1 | 4 | 4 | 3 | 3 |
| ESYT2 | 2 | 2 | 2 | 2 |
| FAF2 | 2 | 2 | 1 | 2 |
| FAR1 | 3 | 3 | 4 | 4 |
| GFPT1 | 1 | 1 | 3 | 3 |
| HLA-A | 2 | 2 | 1 | 1 |
| HSPA5 | 1 | 1 | 3 | 3 |
| LPCAT1 | 1 | 1 | 2 | 2 |
| MAGT1 | 2 | 2 | 2 | 2 |
| MLEC | 2 | 2 | 2 | 2 |
| NCLN | 5 | 5 | 10 | 10 |
| NCSTN | 4 | 4 | 3 | 3 |
| NOMO1 | 9 | 10 | 9 | 10 |
| OS9 | 8 | 8 | 4 | 5 |
| PDIA3 | 4 | 4 | 1 | 1 |
| PSMD2 | 3 | 3 | 3 | 3 |
| PTPLAD1 | 2 | 2 | 2 | 2 |
| PTPN1 | 3 | 3 | 3 | 3 |
| RCN3 | 2 | 2 | 1 | 1 |
| RPN1 | 12 | 13 | 11 | 12 |
| RPTOR | 2 | 2 | 3 | 3 |
| SDF4 | 2 | 2 | 2 | 2 |
| SEC61A1 | 4 | 4 | 5 | 5 |
| SEL1L | 6 | 6 | 2 | 2 |
| SFXN3 | 4 | 4 | 7 | 7 |
| SIL1 | 2 | 2 | 5 | 5 |
| SNX8 | 3 | 3 | 2 | 2 |
| SPTLC1 | 2 | 2 | 1 | 1 |
| ST7L | 1 | 1 | 3 | 3 |
| STT3A | 2 | 2 | 6 | 6 |
| STT3B | 5 | 5 | 7 | 7 |
| SURF4 | 6 | 6 | 4 | 4 |
| TMEM131 | 1 | 1 | 3 | 3 |
| TMEM59 | 2 | 2 | 3 | 3 |
| TMX4 | 1 | 1 | 2 | 2 |
| TTC13 | 4 | 4 | 4 | 4 |
| UBAP2L | 1 | 1 | 4 | 4 |
| VCP | 9 | 9 | 6 | 6 |

**TABLE S2.**
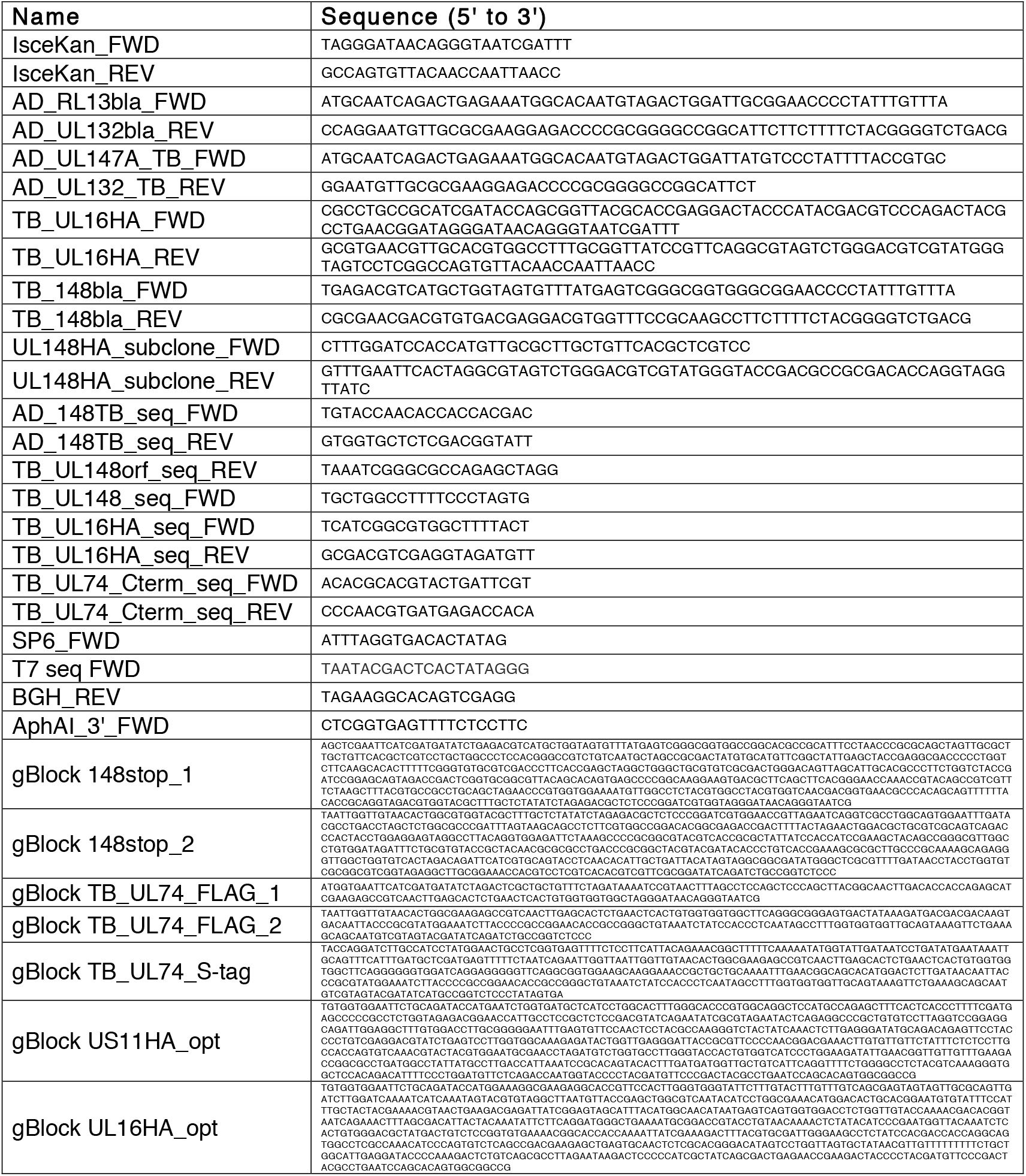
Synthetic DNAs used in the construction and sequence-confirmation of BAC recombinants and plasmids.

**TABLE S3.**
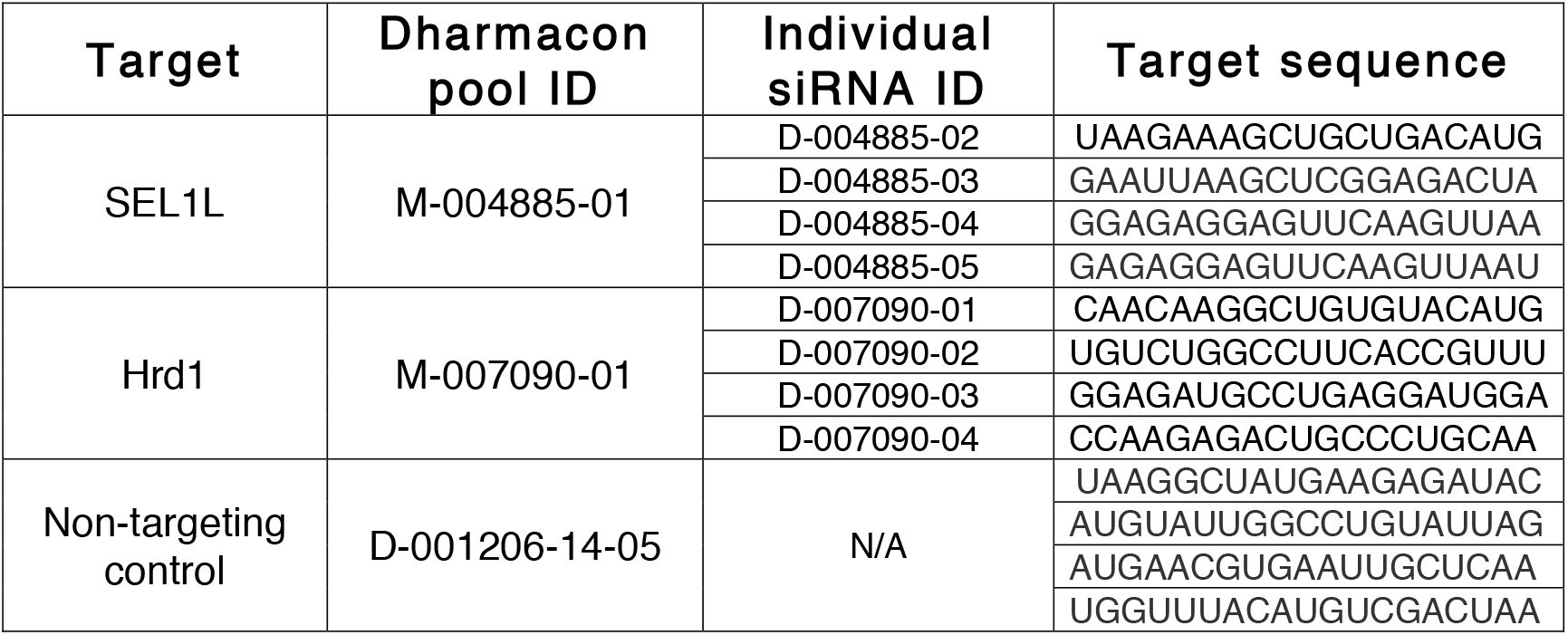
Dharmacon SMARTpool siRNA sequences used in this study.

## SUPPLEMENTAL METHODS

### Construction of recombinant viruses

#### TB40/E expressing HA-tagged UL16

TB40/E expressing HA-tagged UL16 (TB_UL16^HA^) was constructed as follows: A PCR product containing an *ISceI* restriction site upstream of a kanamycin (KAN)- resistance cassette and 40bp flanks homologous to the native UL16 C-terminus was amplified using primers TB_UL16HA_FWD and TB_UL16HA_REV, and TB_148_ISceKan BAC DNA (1) as template. The gel-purified PCR product was electroporated into GS1783 *E. coli* carrying an infectious BAC clone of strain TB40/E BAC, TB40E-BAC4 (2), which was the generous gift of Christian Sinzger (University of Ulm, Germany). After isolating KAN-resistant colonies, the KAN expression cassette was “scarlessly” resolved by inducing expression of the ISceI homing endonuclease and Red recombinase. The final KAN-sensitive TB_UL16^HA^ recombinants were sequence-confirmed using primers TB_UL16HA_seq_FWD and TB_UL16HA_seq_REV.

#### UL 148-null recombinant TB_ 148_STOP_

TB_148bla was first constructed by replacing the *UL 148* ORF of TB_148^HA^ with a beta-lactamase (*bla*) cassette PCR-amplified from plasmid pSP72 (Promega, Inc., Madison, WI) using primers TB_148bla_FWD and TB_148bla_REV. Plasmid pSP72- 148stop-ISceKan was then constructed by Gibson assembly of: (i) gBlock 148stop_1 (ii) gBlock 148stop_2 (iii) ISceKan cassette PCR-amplified from TB_148_ISceKan using primers ISceKan_FWD and ISceKan_REV, and (iv) EcoRV-digested pSP72. The final plasmid was sequence-confirmed using primers SP6_FWD and T7_seq_FWD. pSP72- 148stop-ISceKan was digested by EcoRV to release a 2.1-kb fragment, which was recombined with TB_148bla. Carbenicillin (CARB)-sensitive, KAN-resistant recombinants were resolved of the ISceKan cassette as described above to yield TB_148_STOP_, which was sequence-confirmed using primers TB_UL148_seq_FWD and TB_UL148orf_seq_REV.

#### TB40/E and TB_148_STOP_ expressing S-tagged gO

pSP72-UL74-FLAG-ISceKan was constructed by Gibson Assembly (3) of: (i) EcoRV-digested pSP72, (ii) gBlock TB_UL74_FLAG_1, (iii) gBlock TB_UL74_FLAG_2, and (iv) ISceKan PCR product described above and sequence-confirmed using primers SP6_seq_FWD and T7_seq_FWD. To construct pSP72-UL74-S-ISceKan, pSP72- UL74-FLAG-ISceKan was digested with PpfMI and BglII, and the 3.4-kb fragment was combined with gBlock TB_UL74_S-tag in Gibson assembly to yield pSP72-UL74-S-ISceKan, which was sequence-confirmed using primer AphAI_3’_FWD. pSP72-UL74-S-ISceKan was digested with EcoRV, and the 1.4-kb fragment recombined with the TB40/E or TB_148_STOP_ BAC. KAN-resistant recombinants were resolved of ISceKan as described above to yield TB_gO-S and TB_148_STOP__gO-S, which were sequence-confirmed using primers TB_UL74_Cterm_FWD and TB_UL74_Cterm_REV.

#### ADr131 expressing UL148^HA^ from TB40/E

The vestigial AD169rv *UL148* locus was replaced with *bla* cassette PCR-amplified from template TB_148bla with primers AD_RL13bla_FWD and AD_UL132bla_REV as described above. The ISceKan-disrupted *UL148^HA^* locus from TB_148_ISceKan (1) was PCR-amplified using primers AD_147A_TB_FWD and AD_UL132_TB_REV, recombined in GS1783 *E. coli*with ADr131_148bla, and resolved to yield ADr131_148, which was sequence-confirmed using primers AD_148TB_seq_FWD and AD_148TB_seq_REV.

### Plasmid construction

*pLenti PGK Neo hTERT*. pLenti PGK Neo hTERT was constructed as follows. The *hTERT* ORF was released from pBABE Hygro hTERT (Addgene #1773) by EcoRI/SalI digestion and ligated to XhoI/SalI-digested pENTR1A-no-ccDB (Addgene #17398) to yield pENTR1A hTERT. The *hTERT* ORF was transferred by Gateway cloning to pLenti PGK Neo DEST (Addgene #19067) using LR Clonase II (ThermoFisher #11791-020), resulting in pLenti PGK Neo hTERT.

*UL148^HA^, US11^HA^, and UL16^HA^ expression plasmids*. pEF1α UL148^HA^ was constructed as follows. The *UL148^HA^* ORF was amplified from TB_148^HA^ with primers UL148HA_subclone_FWD and UL148HA_subclone_REV, digested with BamHI /EcoRI, and ligated into pEF1α V5 His C (Invitrogen). pEF1α US11^HA^ and pEF1α UL16^HA^ were constructed by Gibson assembly of EcoRV-digested pEF1α V5 His C (Invitrogen) with gBlock US11 HA_opt or gBlock UL16HA_opt, respectively. Primers T7_seq_FWD and BGH_REV were used to sequence-confirm these constructs.

### hTERT lentivirus production and transduction

5 × 10^5^ 293T cells were co-transfected with pLenti PGK Neo hTERT, psPAX2 (Addgene #12260), and pMD2.G (Addgene #12259) using Mirus Transit-293 reagent according to the manufacturer’s instructions. Lentiviral supernatant was collected at 2 and 3 d post-transfection, 0.45 μm-filtered, supplemented with polybrene (8 μg/mL), and added to subconfluent HFFs. The next day, inocula were removed and cells were washed three times with DPBS. Starting at 4 d post-transduction, cells were serially-passaged in medium containing 500 μg/mL G418. The G418-resistant transduced HFFs that continued to undergo population doublings after control cells senesced were designated hTERT-immortalized HFFs.

### Processing of samples for mass spectrometry analysis

Bands were cut into roughly one cubic millimeter pieces and subjected to an ingel trypsin digest(4). In brief, gel pieces were dehydrated by soaking in acetonitrile for 10 mins, followed by acetonitrile removal and drying by speed-vac. Gel pieces were rehydrated by addition of 50 mM ammonium bicarbonate containing 12.5 ng/μL sequencing-grade trypsin (Promega) for 45 min at 4°C. Excess trypsin solution was removed, and ammonium bicarbonate solution was added to barely cover the gel pieces for overnight incubation at 37°C. Peptides were extracted with 50% acetonitrile/1% formic acid solution, and extracts were dried for 1 hr in a speed-vac. Dried samples were stored at 4°C until analysis. Just before analysis, samples were reconstituted in 510 μL of 2.5% acetonitrile/0.1% formic acid solution (Solvent A).

For preparation of a nano-scale reverse-phase HPLC capillary column, a flame-drawn tip was used to pack 2.6 μm C18 silica beads into a 100 μm (inner diameter) × 30 cm fused silica capillary (5). The column was equilibrated with Solvent A, and a Famos auto sampler (LC Packings, San Francisco, CA) loaded each sample onto the column to allow formation of a gradient. Peptides were then eluted with increasing concentrations of 97.5% acetonitrile/0.1% formic acid.

## REFERENCES

1. Sinzger C, Digel M, Jahn G. 2008. Cytomegalovirus cell tropism. Curr Top Microbiol Immunol 325:63–83.

2. Mocarski ES, Shenk T, Griffiths PD, Pass RF. 2013. Cytomegaloviruses, p 1960–2014. *In* Knipe DM, Howley PM (ed), Fields’ Virology, 6th ed, vol 2. Wolters Kluwer Health/Lippincott Williams & Wilkins, Philadelphia.

3. Goodrum FD, Jordan CT, High K, Shenk T. 2002. Human cytomegalovirus gene expression during infection of primary hematopoietic progenitor cells: a model for latency. Proc Natl Acad Sci U S A 99:16255–60.

4. Maciejewski JP, Bruening EE, Donahue RE, Mocarski ES, Young NS, St Jeor SC. 1992. Infection of hematopoietic progenitor cells by human cytomegalovirus. Blood 80:170–8.

5. Smith MS, Bentz GL, Alexander JS, Yurochko AD. 2004. Human cytomegalovirus induces monocyte differentiation and migration as a strategy for dissemination and persistence. J Virol 78:4444–53.

6. Cannon MJ, Hyde TB, Schmid DS. 2011. Review of cytomegalovirus shedding in bodily fluids and relevance to congenital cytomegalovirus infection. Rev Med Virol 21:240–55.

7. Li G, Kamil JP. 2015. Viral Regulation of Cell Tropism in Human Cytomegalovirus. J Virol 90:626–9.

8. Wang D, Shenk T. 2005. Human cytomegalovirus UL131 open reading frame is required for epithelial cell tropism. J Virol 79:10330–8.

9. Wang D, Shenk T. 2005. Human cytomegalovirus virion protein complex required for epithelial and endothelial cell tropism. Proc Natl Acad Sci U S A 102:18153–8.

10. Hahn G, Revello MG, Patrone M, Percivalle E, Campanini G, Sarasini A, Wagner M, Gallina A, Milanesi G, Koszinowski U, Baldanti F, Gerna G. 2004. Human cytomegalovirus UL131-128 genes are indispensable for virus growth in endothelial cells and virus transfer to leukocytes. J Virol 78:10023–33.

11. Murrell I, Bedford C, Ladell K, Miners KL, Price DA, Tomasec P, Wilkinson GWG, Stanton RJ. 2017. The pentameric complex drives immunologically covert cell-cell transmission of wild-type human cytomegalovirus. Proc Natl Acad Sci U S A 114:6104–6109.

12. Jiang XJ, Adler B, Sampaio KL, Digel M, Jahn G, Ettischer N, Stierhof YD, Scrivano L, Koszinowski U, Mach M, Sinzger C. 2008. UL74 of human cytomegalovirus contributes to virus release by promoting secondary envelopment of virions. J Virol 82:2802–12.

13. Wille PT, Knoche AJ, Nelson JA, Jarvis MA, Johnson DC. 2010. A human cytomegalovirus gO-null mutant fails to incorporate gH/gL into the virion envelope and is unable to enter fibroblasts and epithelial and endothelial cells. J Virol 84:2585–96.

14. Kabanova A, Marcandalli J, Zhou T, Bianchi S, Baxa U, Tsybovsky Y, Lilleri D, Silacci-Fregni C, Foglierini M, Fernandez-Rodriguez BM, Druz A, Zhang B, Geiger R, Pagani M, Sallusto F, Kwong PD, Corti D, Lanzavecchia A, Perez L. 2016. Platelet-derived growth factor-alpha receptor is the cellular receptor for human cytomegalovirus gHgLgO trimer. Nat Microbiol 1:16082.

15. Wu Y, Prager A, Boos S, Resch M, Brizic I, Mach M, Wildner S, Scrivano L, Adler B. 2017. Human cytomegalovirus glycoprotein complex gH/gL/gO uses PDGFR-alpha as a key for entry. PLoS Pathog 13:e1006281.

16. Stegmann C, Hochdorfer D, Lieber D, Subramanian N, Stohr D, Laib Sampaio K, Sinzger C. 2017. A derivative of platelet-derived growth factor receptor alpha binds to the trimer of human cytomegalovirus and inhibits entry into fibroblasts and endothelial cells. PLoS Pathog 13:e1006273.

17. Lemmermann NA, Krmpotic A, Podlech J, Brizic I, Prager A, Adler H, Karbach A, Wu Y, Jonjic S, Reddehase MJ, Adler B. 2015. Non-redundant and redundant roles of cytomegalovirus gH/gL complexes in host organ entry and intra-tissue spread. PLoS Pathog 11:e1004640.

18. Patrone M, Secchi M, Fiorina L, Ierardi M, Milanesi G, Gallina A. 2005. Human cytomegalovirus UL130 protein promotes endothelial cell infection through a producer cell modification of the virion. J Virol 79:8361–73.

19. Scrivano L, Sinzger C, Nitschko H, Koszinowski UH, Adler B. 2011. HCMV spread and cell tropism are determined by distinct virus populations. PLoS Pathog 7:e1001256.

20. Li G, Nguyen CC, Ryckman BJ, Britt WJ, Kamil JP. 2015. A viral regulator of glycoprotein complexes contributes to human cytomegalovirus cell tropism. Proc Natl Acad Sci U S A 112:4471–6.

21. Zhou M, Yu Q, Wechsler A, Ryckman BJ. 2013. Comparative analysis of gO isoforms reveals that strains of human cytomegalovirus differ in the ratio of gH/gL/gO and gH/gL/UL128-131 in the virion envelope. J Virol 87:9680–90.

22. Zhou M, Lanchy JM, Ryckman BJ. 2015. Human Cytomegalovirus gH/gL/gO Promotes the Fusion Step of Entry into All Cell Types, whereas gH/gL/UL128-131 Broadens Virus Tropism through a Distinct Mechanism. J Virol 89:8999–9009.

23. Dunn C, Chalupny NJ, Sutherland CL, Dosch S, Sivakumar PV, Johnson DC, Cosman D. 2003. Human cytomegalovirus glycoprotein UL16 causes intracellular sequestration of NKG2D ligands, protecting against natural killer cell cytotoxicity. J Exp Med 197:1427–39.

24. van de Weijer ML, Bassik MC, Luteijn RD, Voorburg CM, Lohuis MA, Kremmer E, Hoeben RC, LeProust EM, Chen S, Hoelen H, Ressing ME, Patena W, Weissman JS, McManus MT, Wiertz EJ, Lebbink RJ. 2014. A high-coverage shRNA screen identifies TMEM129 as an E3 ligase involved in ER-associated protein degradation. Nat Commun 5:3832.

25. Brodsky JL. 2012. Cleaning up: ER-associated degradation to the rescue. Cell 151:1163–7.

26. Olzmann JA, Kopito RR, Christianson JC. 2013. The mammalian endoplasmic reticulum-associated degradation system. Cold Spring Harb Perspect Biol 5.

27. Christianson JC, Shaler TA, Tyler RE, Kopito RR. 2008. OS-9 and GRP94 deliver mutant alpha1-antitrypsin to the Hrd1-SEL1L ubiquitin ligase complex for ERAD. Nat Cell Biol 10:272–82.

28. Lilley BN, Ploegh HL. 2005. Multiprotein complexes that link dislocation, ubiquitination, and extraction of misfolded proteins from the endoplasmic reticulum membrane. Proc Natl Acad Sci U S A 102:14296–301.

29. Mueller B, Lilley BN, Ploegh HL. 2006. SEL1L, the homologue of yeast Hrd3p, is involved in protein dislocation from the mammalian ER. J Cell Biol 175:261–70.

30. Lilley BN, Ploegh HL. 2004. A membrane protein required for dislocation of misfolded proteins from the ER. Nature 429:834–40.

31. Alcock F, Swanton E. 2009. Mammalian OS-9 is upregulated in response to endoplasmic reticulum stress and facilitates ubiquitination of misfolded glycoproteins. J Mol Biol 385:1032–42.

32. Cha TA, Tom E, Kemble GW, Duke GM, Mocarski ES, Spaete RR. 1996. Human cytomegalovirus clinical isolates carry at least 19 genes not found in laboratory strains. J Virol 70:78–83.

33. Elbein AD, Tropea JE, Mitchell M, Kaushal GP. 1990. Kifunensine, a potent inhibitor of the glycoprotein processing mannosidase I. J Biol Chem 265:15599–605.

34. Bernasconi R, Galli C, Calanca V, Nakajima T, Molinari M. 2010. Stringent requirement for HRD1, SEL1L, and OS-9/XTP3-B for disposal of ERAD-LS substrates. J Cell Biol 188:223–35.

35. Hosokawa N, Kamiya Y, Kamiya D, Kato K, Nagata K. 2009. Human OS-9, a lectin required for glycoprotein endoplasmic reticulum-associated degradation, recognizes mannose-trimmed N-glycans. J Biol Chem 284:17061–8.

36. Ron E, Shenkman M, Groisman B, Izenshtein Y, Leitman J, Lederkremer GZ. 2011. Bypass of glycan-dependent glycoprotein delivery to ERAD by up-regulated EDEM1. Mol Biol Cell 22:3945–54.

37. Luganini A, Cavaletto N, Raimondo S, Geuna S, Gribaudo G. 2017. Loss of the Human Cytomegalovirus US16 Protein Abrogates Virus Entry into Endothelial and Epithelial Cells by Reducing the Virion Content of the Pentamer. J Virol 91.

38. Jiang XJ, Sampaio KL, Ettischer N, Stierhof YD, Jahn G, Kropff B, Mach M, Sinzger C. 2011. UL74 of human cytomegalovirus reduces the inhibitory effect of gH-specific and gB-specific antibodies. Arch Virol 156:2145–55.

39. Wei X, Decker JM, Wang S, Hui H, Kappes JC, Wu X, Salazar-Gonzalez JF, Salazar MG, Kilby JM, Saag MS, Komarova NL, Nowak MA, Hahn BH, Kwong PD, Shaw GM. 2003. Antibody neutralization and escape by HIV-1. Nature 422:307–12.

40. Huber MT, Compton T. 1999. Intracellular formation and processing of the heterotrimeric gH-gL-gO (gCIII) glycoprotein envelope complex of human cytomegalovirus. J Virol 73:3886–92.

41. Yang M, Omura S, Bonifacino JS, Weissman AM. 1998. Novel aspects of degradation of T cell receptor subunits from the endoplasmic reticulum (ER) in T cells: importance of oligosaccharide processing, ubiquitination, and proteasome-dependent removal from ER membranes. J Exp Med 187:835–46.

42. Bonifacino JS, Suzuki CK, Lippincott-Schwartz J, Weissman AM, Klausner RD. 1989. Pre-Golgi degradation of newly synthesized T-cell antigen receptor chains: intrinsic sensitivity and the role of subunit assembly. J Cell Biol 109:73–83.

43. Chandramouli S, Malito E, Nguyen T, Luisi K, Donnarumma D, Xing Y, Norais N, Yu D, Carfi A. 2017. Structural basis for potent antibody-mediated neutralization of human cytomegalovirus. Sci Immunol 2.

44. Ciferri C, Chandramouli S, Donnarumma D, Nikitin PA, Cianfrocco MA, Gerrein R, Feire AL, Barnett SW, Lilja AE, Rappuoli R, Norais N, Settembre EC, Carfi A. 2015. Structural and biochemical studies of HCMV gH/gL/gO and Pentamer reveal mutually exclusive cell entry complexes. Proc Natl Acad Sci U S A 112:1767–72.

45. Ryckman BJ, Rainish BL, Chase MC, Borton JA, Nelson JA, Jarvis MA, Johnson DC. 2008. Characterization of the human cytomegalovirus gH/gL/UL128-131 complex that mediates entry into epithelial and endothelial cells. Journal of Virology 82:60-70.

46. Stanton RJ, Baluchova K, Dargan DJ, Cunningham C, Sheehy O, Seirafian S, McSharry BP, Neale ML, Davies JA, Tomasec P, Davison AJ, Wilkinson GW. 2010. Reconstruction of the complete human cytomegalovirus genome in a BAC reveals RL13 to be a potent inhibitor of replication. J Clin Invest 120:3191–208.

47. Zhang L, Zhou M, Stanton R, Kamil JP, Ryckman BJ. 2018. Expression levels of glycoprotein O (gO) vary between strains of human cytomegalovirus, influencing the assembly of gH/gL complexes and virion infectivity. bioRxiv doi:10.1101/299222.

48. Wang D, Li G, Schauflinger M, Nguyen CC, Hall ED, Yurochko AD, von Einem J, Kamil JP. 2013. The ULb’ region of the human cytomegalovirus genome confers an increased requirement for the viral protein kinase UL97. J Virol 87:6359–76.

49. Kärber G. 1931. Beitrag zur kollektiven Behandlung pharmakologischer Reihenversuche. Archiv f experiment Pathol u Pharmakol 162: 480–483.

50. Spearman C. 1908. The Method of “Right and Wrong Cases” (Constant Stimuli) without Gauss’s Formula. Br J Psychol 2:227–242

51. Urban M, Britt W, Mach M. 1992. The dominant linear neutralizing antibody-binding site of glycoprotein gp86 of human cytomegalovirus is strain specific. J Virol 66:1303–11.

52. Spaete RR, Saxena A, Scott PI, Song GJ, Probert WS, Britt WJ, Gibson W, Rasmussen L, Pachl C. 1990. Sequence requirements for proteolytic processing of glycoprotein B of human cytomegalovirus strain Towne. J Virol 64:2922–31.

53. Sanchez V, Greis KD, Sztul E, Britt WJ. 2000. Accumulation of virion tegument and envelope proteins in a stable cytoplasmic compartment during human cytomegalovirus replication: characterization of a potential site of virus assembly. J Virol 74:975–86.

54. Sinzger C, Hahn G, Digel M, Katona R, Sampaio KL, Messerle M, Hengel H, Koszinowski U, Brune W, Adler B. 2008. Cloning and sequencing of a highly productive, endotheliotropic virus strain derived from human cytomegalovirus TB40/E. J Gen Virol 89:359–68.

55. Hobom U, Brune W, Messerle M, Hahn G, Koszinowski UH. 2000. Fast screening procedures for random transposon libraries of cloned herpesvirus genomes: mutational analysis of human cytomegalovirus envelope glycoprotein genes. J Virol 74:7720–9.

56. Li G, Rak M, Nguyen CC, Umashankar M, Goodrum FD, Kamil JP. 2014. An epistatic relationship between the viral protein kinase UL97 and the UL133-UL138 latency locus during the human cytomegalovirus lytic cycle. J Virol 88:6047–60.

57. Tischer BK, Smith GA, Osterrieder N. 2010. En passant mutagenesis: a two step markerless red recombination system. Methods Mol Biol 634:421–30.

58. Tischer BK, von Einem J, Kaufer B, Osterrieder N. 2006. Two-step red-mediated recombination for versatile high-efficiency markerless DNA manipulation in Escherichia coli. Biotechniques 40:191–7.

59. Richards FM, Vithayathil PJ. 1959. The preparation of subtilisn-modified ribonuclease and the separation of the peptide and protein components. J Biol Chem 234:1459–65.

60. Eng JK, McCormack AL, Yates JR. 1994. An approach to correlate tandem mass spectral data of peptides with amino acid sequences in a protein database. J Am Soc Mass Spectrom 5:976–89.

## SUPPLEMENTAL REFERENCES

1. Li G, Nguyen CC, Ryckman BJ, Britt WJ, Kamil JP. 2015. A viral regulator of glycoprotein complexes contributes to human cytomegalovirus cell tropism. Proc Natl Acad Sci U S A 112:4471–6.

2. Sinzger C, Hahn G, Digel M, Katona R, Sampaio KL, Messerle M, Hengel H, Koszinowski U, Brune W, Adler B. 2008. Cloning and sequencing of a highly productive, endotheliotropic virus strain derived from human cytomegalovirus TB40/E. J Gen Virol 89:359–68.

3. Gibson DG, Young L, Chuang RY, Venter JC, Hutchison CA, 3rd, Smith HO. 2009. Enzymatic assembly of DNA molecules up to several hundred kilobases. Nat Methods 6:343–5.

4. Shevchenko A, Jensen ON, Podtelejnikov AV, Sagliocco F, Wilm M, Vorm O, Mortensen P, Shevchenko A, Boucherie H, Mann M. 1996. Linking genome and proteome by mass spectrometry: large-scale identification of yeast proteins from two dimensional gels. Proc Natl Acad Sci U S A 93:14440–5.

5. Peng J, Gygi SP. 2001. Proteomics: the move to mixtures. J Mass Spectrom 36:1083–91.

6. Gray KA, Yates B, Seal RL, Wright MW, Bruford EA. 2015. Genenames.org: the HGNC resources in 2015. Nucleic Acids Res 43:D1079–85.

